# Drone-derived canopy height predicts biomass across non-forest ecosystems globally

**DOI:** 10.1101/2020.07.16.206011

**Authors:** A.M. Cunliffe, K. Anderson, F. Boschetti, R.E. Brazier, H.A. Graham, I.H. Myers-Smith, T. Astor, M.M. Boer, L. Calvo, P.E. Clark, M.D. Cramer, M.S. Encinas-Lara, S.M. Escarzaga, J.M. Fernández-Guisuraga, A.G. Fisher, K. Gdulová, B.M. Gillespie, A. Griebel, N.P. Hanan, M.S. Hanggito, S. Haselberger, C.A. Havrilla, P. Heilman, W. Ji, J.W. Karl, M. Kirchhoff, S. Kraushaar, M.B. Lyons, I. Marzolff, M.E. Mauritz, C.D. McIntire, D. Metzen, L.A. Méndez-Barroso, S.C. Power, J. Prošek, E. Sanz-Ablanedo, K.J. Sauer, D. Schulze-Brüninghoff, P. Šímová, S. Sitch, J.L. Smit, C.M. Steele, S. Suárez-Seoane, S.A. Vargas, M.L. Villarreal, F. Visser, M. Wachendorf, H. Wirnsberger, R. Wojcikiewicz

## Abstract

Non-forest ecosystems, dominated by shrubs, grasses and herbaceous plants, provide ecosystem services including carbon sequestration and forage for grazing, yet are highly sensitive to climatic changes. Yet these ecosystems are poorly represented in remotely-sensed biomass products and are undersampled by *in-situ* monitoring. Current global change threats emphasise the need for new tools to capture biomass change in non-forest ecosystems at appropriate scales. Here we assess whether canopy height inferred from drone photogrammetry allows the estimation of aboveground biomass (AGB) across low-stature plant species sampled through a global site network. We found mean canopy height is strongly predictive of AGB across species, demonstrating standardised photogrammetric approaches are generalisable across growth forms and environmental settings. Biomass per-unit-of-height was similar *within*, but different *among*, plant functional types. We find drone-based photogrammetry allows for monitoring of AGB across large spatial extents and can advance understanding of understudied and vulnerable non-forested ecosystems across the globe.

## Introduction

Non-forest ecosystems, dominated by shrub and herbaceous plants, cover about 70% of the Earth’s land surface^1^ and account for around 35% of all aboveground biomass (AGB)^2^. They provide multiple ecosystem services, playing dominant roles in the long-term trends and interannual variability of the global carbon cycle^3,4^, and are highly significant for grazing and agriculture^5^. Grassland, shrubland, Arctic tundra, savanna and proglacial montane landscapes are often more sensitive and respond faster to changes in climate than forests^6^, but have received less systematic research attention^1,7,8^. Non-destructive measurements of canopy height and biomass are fundamental requirements for plant science^9–11^ to understand the roles of these ecosystems in climate change mitigation, sustainable food production and land management^12–14^. However, measuring biomass with *in situ* measurements is labour intensive and thus prone to undersampling, particularly in ecosystems that are spatially heterogeneous and/or temporally dynamic, putting on (and losing) biomass rapidly^1,15–17^. Gaps in available data mean that biomass dynamics are not being captured in many important ecosystems across the globe, hindering the calibration and validation of vegetation models and products derived from satellite observations^7,14^. The lack of accurate biomass data limits our ability to track changes and predict future responses in globally relevant non-forest ecosystems.

Improving the accuracy of biomass data in non-forest biomes requires approaches that are: (i) sensitive to small differences in AGB, (ii) sufficiently inexpensive to be adopted worldwide, and allow (iii) spatially continuous sampling across (iv) representative areas at (v) temporal frequencies appropriate for dynamic ecosystems^14,16^. The most accurate non-destructive estimates of AGB are generally obtained from *in situ* measurements of attributes such as plant cover, height and stem diameters, using allometric functions fitted to harvested biomass observations. Canopy volume, the product of height and cover, is often the strongest predictor of AGB for shrubs, herbs and other low-stature plants^15,18–22^. Remote-sensing approaches have been widely used to extend the coverage of biomass predictions. Biomass can be predicted from airborne LiDAR (Light Detection and Ranging) in shrublands and savannas^23^, but the footprints sampled by LiDAR can be insensitive to fine-scale changes in plant structure and these data are expensive and unavailable in many areas. Biomass estimates computed from spectral reflectance are often highly uncertain due to asymptotic relationships between AGB and surface reflectance and variable soil albedo^6,22^. Globally-available biomass products from space-based sensors such as LiDAR, synthetic-aperture radar or vegetation optical depth are either insensitive and/or poorly calibrated and validated in low-biomass (<20 Mg ha^-1^) ecosystems^1,7,8,14,24^.

Photogrammetry using aerial images acquired with unmanned aerial systems (UAS, herein ‘drones’) could greatly improve quantification of AGB in non-forest ecosystems, both directly at a local scale and indirectly by improving the calibration and validation of biomass products obtained from coarser scale remotely-sensed observations. Advances in photogrammetry, particularly structure-from-motion (SfM) with multi-view stereopsis^25^, have made it possible to capture 3D representations of plants, quantitatively describing their fine-scale structure with an unprecedented level of detail^26,27^. SfM allows objective measurements of canopy height at sub-decimetre spatial grain for a wide range of plant growth forms^18–20,27–31^. Lightweight and inexpensive drones enable vegetation sampling at temporal intervals appropriate for highly dynamic ecosystems^16,32^ and can be used for detailed surveys over extents of 1-10 ha, covering more representative areas of heterogeneous ecosystems^15,27^ that allow spatially explicit comparison with other biomass estimates^1,14,32^.

Fully realising the potential of drone photogrammetry in plant science requires reproducible workflows, which minimise biases^30,33,34^. Over the past few years, thousands of hectares of low-stature ecosystems have been surveyed with drones across the globe, yielding information-rich datasets. However, drone-photogrammetry products are sensitive to the ways in which data are (i) collected (e.g., ground sampling distance, image overlap, viewing geometry, spatial control, illumination conditions)^27,30,34–38^, (ii) processed (e.g., software, lens model, specification of control accuracy, selection of processing quality, depth filtering)^27,36–38^, and (iii) analysed (e.g., canopy height metrics, spatial grain and interpolation, statistical treatment)^20,27–29,31^. These sensitivities are more pronounced for subjects with complex texture, such as vegetation, and hinder comparisons between products obtained from different workflows. To maximise the value of these approaches, standardised and reproducible protocols are needed, but few efforts currently exist to advance this aim. Addressing critical knowledge gaps in plant science with drone photogrammetry requires knowledge of the relationships between photogrammetrically derived canopy height and AGB across the range of plants and ecosystems in which they will be applied, and systematic understanding of the possible influences of environmental conditions (e.g. wind speed and illumination)^19,27,30,35,39^. In this study, we apply a new, standardised approach for airborne allometric inference of biomass for non-forested ecosystems globally using drone photogrammetry. We asked the following research questions: (1) Does canopy height derived from drone photogrammetry correspond with AGB at the species-level? (2) Does photogrammetry-derived canopy height correspond with AGB at the PFT-level? (3) Are relationships between reconstructed canopy height and biomass influenced by wind speed and (4) and solar elevation?

Using rigorous, consistent protocols^33^, we conducted a novel, globally coordinated experiment to sample 36 sites, encompassing a diverse range of non-forest ecosystems, including semi-arid and temperate grasslands and shrublands, Arctic tundra, savanna and proglacial montane sites (Fig. 1B), spanning from 71° North to 37° South, across North America, Europe, Australia and Africa (Fig. 1A). Our study includes photogrammetric reconstructions from 38 different surveys (Supplementary Table 1), sampling 50 low-stature plant species across six PFTs including ferns, forbs, graminoids, shrubs, succulents and trees that cover phylogenetic diversity including non-flowering plants and the most species-rich clades of flowering plants (including monocots and eudicots). To calibrate our allometric models, we sampled 741 harvest plots, with AGB ranging from 9 g m^-2^ to 7,892 g m^-2^ and mean (maximum) canopy heights ranging from 0 m to 1.9 m (0.01 m to 6.7 m). Our sample achieved a more than twenty-fold improvement in the coverage of harvest plots, species and sites compared to previous photogrammetry vegetation studies (Fig. 1C)^20,28,29^. We fitted plant functional type (PFT) and species-specific models that predict AGB from fine-grained canopy height as determined by SfM photogrammetry. Mean canopy height, sampled at fine (centimetre) spatial grain, integrated canopy cover and height as well as foliage density. The consideration of these multiple plant size attributes is key to robust prediction of biomass.

**Fig. 1.**
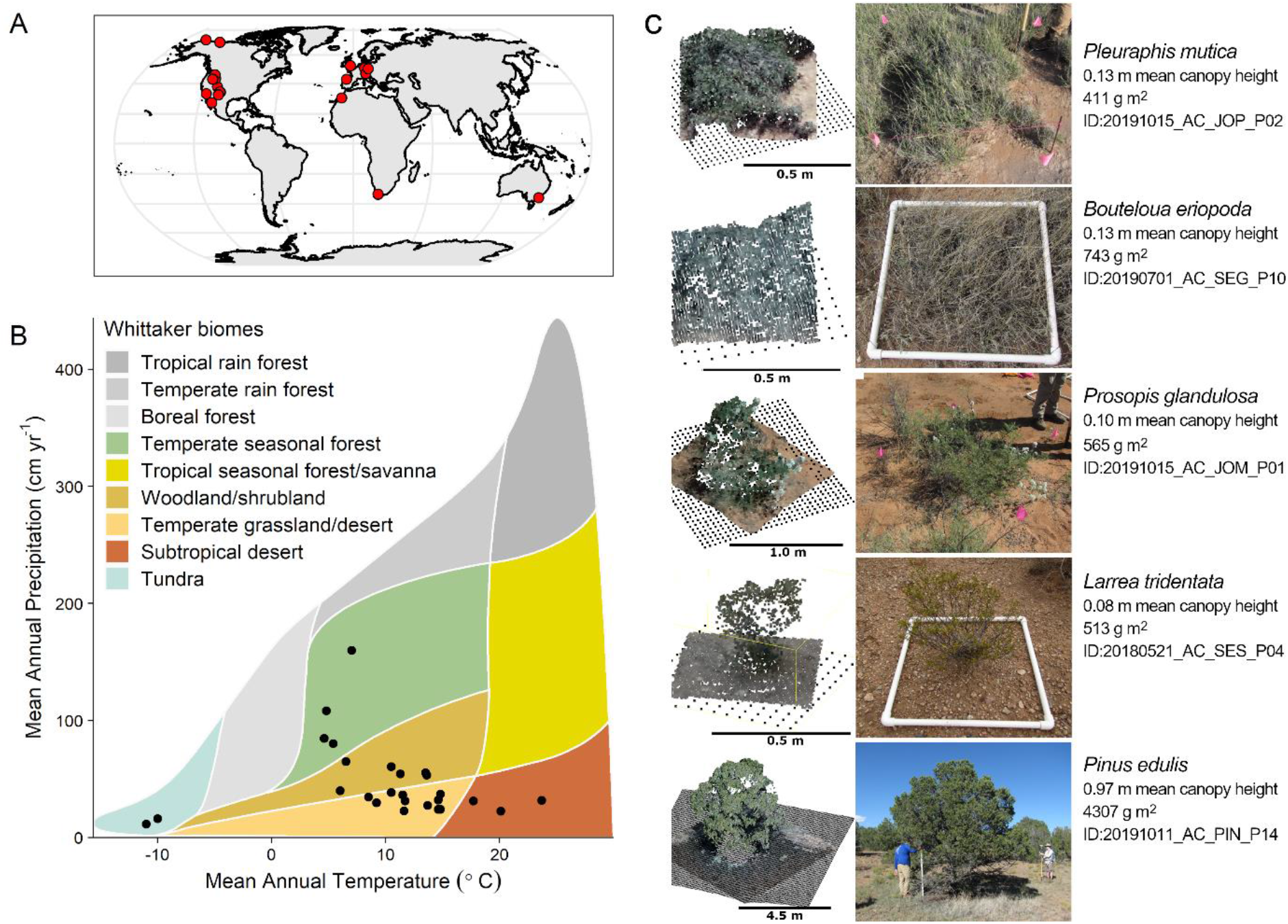
Point clouds derived from drone surveys consistently provided structural reconstructions of plants across non-forested ecosystems. **A** depicts the geographic distribution of our sites, spanning four continents. **B** depicts the bioclimatic distribution of sites in terms of annual average precipitation and temperature. We sampled five biomes where low-stature vegetation is often dominant, representing every appropriate (non-forest) biome described by Whitaker^40^. **C** Reconstructed point clouds corresponded well with photographs of harvest plots. The grid of black points represents the underlying terrain model.

## Methods

### Site and Species selection

We focused our efforts on low-stature phenotypes in non-forest ecosystems, including grasslands, shrublands with open and closed canopies, and woody savannas. Low-stature ecosystems are understudied and tools for quantifying forest biomass are better represented in the existing literature^1,7,8^. While photogrammetry can be used to characterise forest canopies^30,31,35^, we consider forest ecosystems better candidates for observation with active remote sensing approaches such as synthetic-aperture radar^7^, vegetation optical depth and LiDAR^52^. We selected species that were regionally widespread, accessible and would inform ongoing research efforts, but excluded extensively modified vegetation such as managed hedges. Sampling was undertaken during seasonal peak canopy cover to minimise differences arising from phenophase, although plant development and allometric relationships may still vary especially in more water-limited ecosystems^41^. The data collection protocol was comprehensively described by Cunliffe and Anderson^33^. Two study sites (‘SES’ and ‘SEG’) were sampled on consecutive years, giving 38 surveys from 36 sites (Supplementary Table 1).

### Aerial imaging surveys

Harvest sites were surveyed using drones to acquire aerial red-green-blue images. For each site, two sets of survey flights were undertaken, the first obtaining nadir imagery to attain a spatial grain of ca. 5 mm per pixel at the canopy top, and the second obtaining oblique (ca. 20° from nadir) images from ca. 4 m higher. Survey altitudes varied depending on the resolution and field-of-view of the sensors and the canopy height^30^, but were typically 20 m above the canopy. The different perspectives afforded by the nadir and slightly higher, convergent surveys helps to improve the stability of the camera network^36,37,45,53–56^. Both survey flights obtained 75% forward and side overlap, together capturing at least 30 images for each part of the study area. The high image overlap facilitated tie point matching in the vegetated scenes. Wind speeds were generally recorded using handheld anemometers immediately prior to the survey^47^. Our sampling protocol^33^ was optimised for smaller plants of up to ca. 3 m in height. To support feature matching in texturally complex scenes containing taller vegetation (e.g., mature *Juniperus monosperma* or *Pinus edulis*), higher survey altitudes could be used to help minimise excessive parallax (i.e., excessive scene changes between overlapping images).

A key requirement for photogrammetric surveys is the inclusion of adequate spatial control^38,45^. Our photogrammetric reconstructions used thirteen ground markers, each measuring ca. 20 cm x 20 cm, deployed across each site and geolocated to a typical precision of ± 0.015 m horizontally and ± 0.03 m vertically. Further details on the sites and survey equipment are provided in Supplementary Table 1. Images intended for photogrammetric analysis should ideally not be geometrically corrected in-camera prior to further distortion correction. Such in-camera processing is a problem for JPG-format image files from cameras like the widely used DJI Phantom 4 Advanced/Pro FC6310 camera, and so capturing RAW-format images can help avoid this error source^38,45^. We anticipate ongoing improvements to camera geolocation and orientation information from drone systems will continue to improve the accuracy and reliability of the camera parameter estimation, particularly in densely vegetated and thus texturally complex settings (Supplementary Note 1)^34,38,45,57,58^.

### Vegetation harvests

We used an area-based approach to enable sampling in ecosystems with continuous or coalesced canopies, while also sampling individual plants where these were naturally isolated from other plants^33,59^. We selected harvest plots to sample across the natural range of canopy heights observed at each site, in order to estimate the allometric models more efficiently as well as to test the form of the relationship between mean canopy height and biomass^46^. Plots were chosen to try to ensure that ≥ 90% of the biomass and ≥ 90% of the foliar volume within each plot were from the target species. We aimed for a minimum harvest plot size of 0.5 m x 0.5 m to reduce the possible effects of co-registration errors^22^. The corners of each plot were geolocated with high-precision GNSS before all standing biomass was harvested to ground-level (or the moss level for *Salix richardsonii* and *Arctophila fulva*)^22^. Biomass was then dried at ca. 50-80°C until reaching a constant weight over a 24-hour period. For some of the largest taxa (*Adenostoma fasciculatum, Adenostoma sparsifolium, Atriplex polycarpa, Ericameria nauseosa, Juniperus monosperma, Launaea arborescens, Pinus edulis* and *Prosopis velutina*), freshly harvested biomass was weighed in the field and representative sub-samples were then dried to determine moisture contents^59^. Plot areas were computed from corner coordinates, unless a quadrat was used during harvesting in which case the area of the quadrat was used to minimise propagating errors from GNSS-coordinates.

### Image-based modelling

Aerial images were processed using SfM photogrammetry, using established workflows and following our previous studies^27,59^. Geotagged image data and ground-control marker coordinates were imported into AgiSoft PhotoScan Professional v1.4.3 (now Metashape; http://www.agisoft.com) and converted to UTM coordinate reference systems. Image sharpness was measured using PhotoScan’s image quality tool, all images had an image sharpness score of ≥ 0.5^37^. Tie points were matched and cameras aligned using PhotoScan’s highest quality setting, a key point limit of 40,000, a tie point limit of 8,000, with generic and reference pair preselection enabled, and adaptive camera model fitting disabled. During camera self-calibration we enabled the following lens parameters: Focal length (f), principal point (cx, cy), radial distortion (k1, k2), tangential distortion (p1, p2), aspect ratio and skew coefficient (b1, b2). Most cameras had global shutters but rolling shutter corrections were used when appropriate. Reference parameters were set to: camera location accuracy = XY ± 20 m, Z ± 50 m; marker location accuracy = XY ± 0.02 m, Z ± 0.05 m; marker projection accuracy was set to 2 pixels; tie point accuracy was set to either the mean root mean square reprojection error or one, whichever was greater. The result of camera alignment was a sparse point cloud that was then filtered and points with reprojection error above 0.45 pixels were excluded from further analysis. An operator reviewed the sparse point clouds and estimated camera positions to verify their plausibility. Any obviously erroneous tie points were removed manually. Geolocated markers were placed by an operator on ten projected images for each of the 13 ground control points. Ten of these markers were used to constrain the photogrammetric reconstructions spatially^60^, while the remaining three were used for independent evaluation of each reconstruction. The three markers used for accuracy assessment were deselected before the interior and exterior camera parameters were optimised. Any obviously implausible camera positions were refined after marker placement and optimisation. All cameras were usually aligned and used for multi-view stereopsis (dense point cloud generation), using the ultrahigh quality setting with mild depth filtering to preserve finer details of the vegetation^27,29,30^. For further discussion of some of the limitations of this approach, see Supplementary Note 1. Dense point clouds were exported in the laz format, with point coordinate and RGB attributes.

### Digital terrain models

An essential requirement for deriving canopy height models from photogrammetry-derived point clouds is a digital terrain model, which must be sufficiently accurate and detailed with respect to canopy heights and topographic complexity^31^. We used terrain models interpolated with Delaunay triangulation between the GNSS-observations of the harvest plot corners (Fig. 1C). In instances where plant canopies are discontinuous in space, suitable terrain models may be extracted from the photogrammetric point cloud^20,27^. Other options can include extracting terrain models from photogrammetric drone surveys during leaf-off conditions (or post-harvest, if applicable), LiDAR surveys^61^ or walkover surveys with GNSS instruments.

### Calculation of canopy heights

Point clouds were analysed with PDAL (v2.1.0)^62^. The point cloud representing each harvest plot was subset using the GNSS-observed corner coordinates. In a few instances where plot infrastructure (e.g., marker posts or flags) was visible in the point cloud (n=20 plots), these points were manually assigned to a noise class and excluded from canopy height calculations. Within each plot, the height-above-ground of each point was calculated relative to the terrain model and any points with a negative height-above-ground were set to zero^20,27^. Using a 0.01 m resolution grid, we calculated the maximum point height in each grid cell. For cells containing no points, we interpolated heights using inverse distance weighting considering an array of 7 × 7 cells using a power of one, and cells with no neighbouring points in that area remained empty. Plot-level mean canopy height was then extracted from this grid of local maxima elevations.

### Statistical analysis

Statistical analyses were conducted in R v3.6.1^63^. Sun elevations during each survey were computed with the Astral package^64^. We produced the climate space plot using the plotbiomes R package^65^ based on the biomes described by Whittaker^40^. We excluded 13 bryophyte plots from two rocky sites where we were unable to extract meaningful canopy height observations (Supplementary Fig. 5) and 16 graminoid plots from one grassland site (‘WSP’) that could not be reconstructed (Supplementary Fig. 6, Supplementary Note 1).

We used ordinary least squares regression to fit separate linear models predicting AGB observations from mean canopy height for each PFT and for each species with four or more observations. We considered ferns, forbs, graminoids, shrubs, trees and succulents as PFTs, and constrained the y-intercept to zero in order to ensure zero canopy height predicted zero biomass. Model performance was validated using leave-one-out cross-validation (LOOCV) to compute the mean out-of-sample prediction error, which was divided by the model slope to obtain relative errors for each model^31,66^.

To test the influence of wind speed on allometric functions, we fitted a generalised linear mixed model (GLMM) to predict total biomass as a function of canopy height and wind speed as fixed-effects and PFT as a random-effect based on a gamma error distribution with an identity link function, using the ‘lme4’ package (v1.1-23)^67^ (Supplementary Table 3). Succulents were excluded from this model because their inclusion prevented model convergence, possibly because this PFT had a much steeper slope between height:biomass (Table 1, Fig. 2) and/or because they may be less influenced by wind speed (Supplementary Fig. 2A). To illustrate the effect of wind speed, we used the ‘ggeffects’ pacakge (v0.15)^68^ to simulate the relationship between height and biomass for three levels of wind speed using the GLMM (Fig. 3A), and plotted the slope of biomass-height models (±83% confidence interval^69^) against wind speed at the PFT- (Supplementary Fig. 2A) and species-levels (Supplementary Fig. 3).

**Fig. 2.**
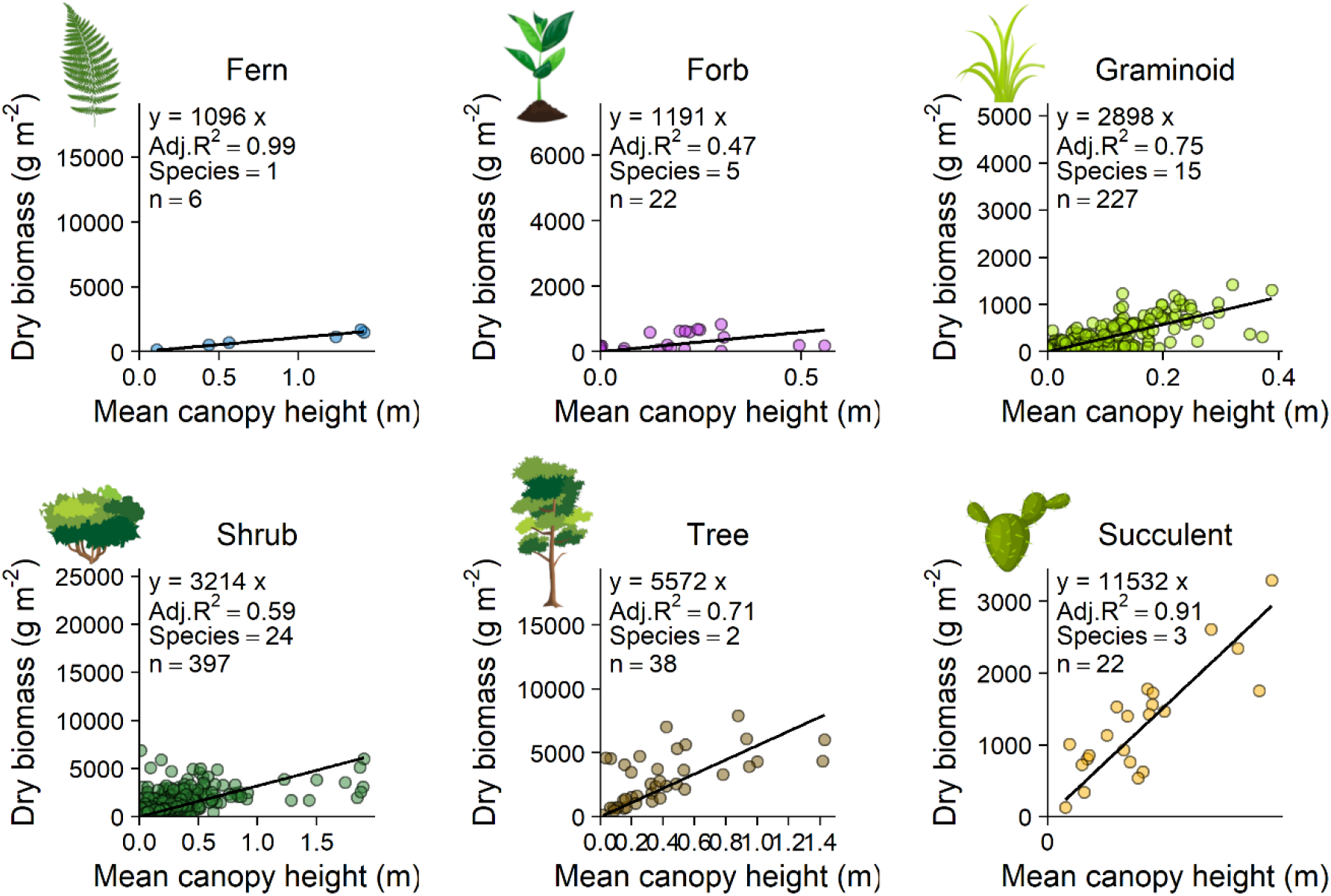
Photogrammetrically derived canopy height was a strong predictor of biomass within plant functional types. A constant X:Y ratio was used for all plots, so model slopes can be compared visually even though axis ranges vary. Model slopes were generally similar within, but differed between, plant functional types. ‘Species’ indicates the number of species pooled for each plant functional type and black lines are linear models with intercepts constrained through the origin. Full model results are included in Table 1.

**Table 1.** Parameters for linear models fitted to each plant functional type. LOOCV is the prediction error from Leave-One-Out Cross-Validation divided by the slope.

| Plant functional type | n | n of surveys | Slope<br>g m<sup>-2</sup> | Residual standard error<br>g m<sup>-2</sup> | Adj. R<sup>2</sup> | t-statistic | P value | LOOCV<br>% |
| --- | --- | --- | --- | --- | --- | --- | --- | --- |
| Fern | 6 | 1 | 1,096 | 53 | 0.99 | 20.558 | <0.0001 | 12.0 |
| Forb | 22 | 3 | 1,191 | 262 | 0.47 | 4.534 | 0.0002 | 19.0 |
| Graminoid | 227 | 17 | 2,898 | 112 | 0.75 | 25.786 | <0.0001 | 3.7 |
| Shrub | 397 | 24 | 3,214 | 134 | 0.59 | 23.823 | <0.0001 | 11.6 |
| Succulent | 22 | 3 | 11,532 | 760 | 0.91 | 15.159 | <0.0001 | 2.6 |
| Tree | 38 | 2 | 5,572 | 577 | 0.71 | 9.654 | <0.0001 | 16.7 |

To test the influence of cloud cover on allometric functions, we fitted a linear mixed model (LMM) to predict total biomass as a function of canopy height, with PFT as a random-effect and cloud cover as fixed-effects, using the ‘lmerTest’ package (v3.1-2)^70^ (Supplementary Table 4). Cloud cover was coded as a binary factor, with relatively clear sky (n=620) and cloudy conditions where the sun was obscured (n=80, sky codes ≥ 6 after^71^, Supplementary Table 6). To illustrate the effect of sun elevation, we simulated the modelled relationship between height and biomass for the two levels of cloud cover using the LMM (Supplementary Fig. 4).

To test the influence of sun elevation on allometric functions, we fitted a LMM to predict total biomass as a function of canopy height and sun elevation as fixed-effects and PFT as a random-effect, using the ‘lmerTest’ package (v3.1-2)^70^ (Supplementary Table 5). We only included observations (n=620) collected under relatively clear sky conditions (sky codes ≤ 5, after^71^) when scene illumination was minimally modulated by clouds. To illustrate the effect of sun elevation, we simulated the modelled relationship between height and biomass for three levels of sun elevation using the LMM (Fig. 3B), and plotted the slope of biomass-height models (±83% confidence interval^69^) against sun elevation at the PFT- (Supplementary Fig. 3B) and species-level (Supplementary Fig. 5). There was insufficient replication to allow convergence of more complex model structures including species nested within PFT or site as random-effects. We evaluated diagnostics for all model visually using the R package ‘performance’ (v0.4.6)^72^.

**Fig. 3.**
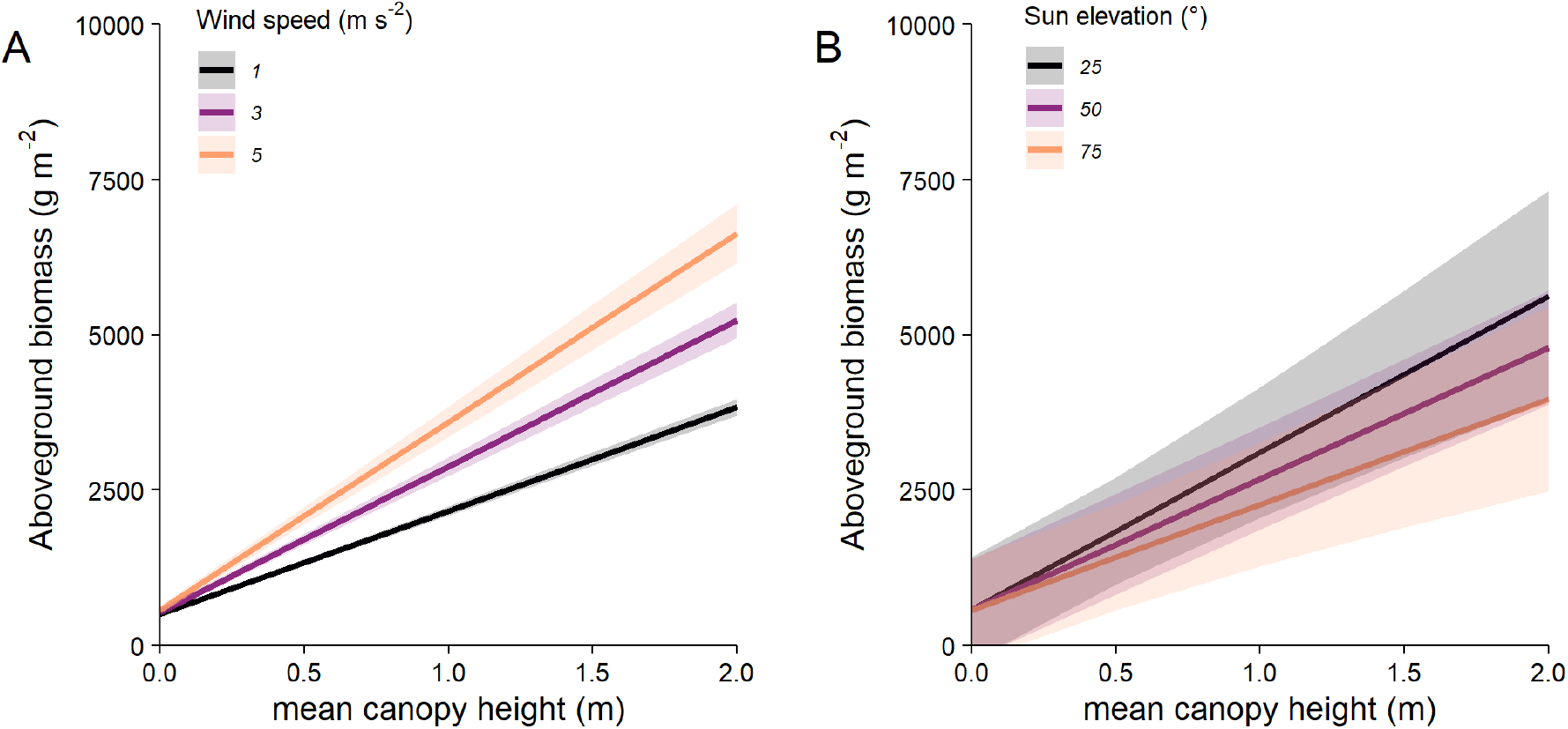
Reconstructed plant height and thus height-biomass relationships were systematically influenced by wind speed but were insensitive to illumination conditions. Mean predicted aboveground biomass variation over the range of observed mean canopy height, estimated for a range of three wind speeds and sun elevations. Wind speed has a statistically clear and positive effect on the relationship between height and biomass (**A**) (Supplementary Figs. 2A and 3, Supplementary Table 3), but sun elevation had no significant effect on the relationship between height and biomass (**B**) (Supplementary Figs. 2B and 5, Supplementary Table 5). Shaded areas represent 95% confidence intervals on the model predictions.

## Results

We found photogrammetrically measured mean canopy height was strongly predictive of AGB at the species-level. Linear models with a zero-intercept provided good approximations of the relationships between mean canopy height and AGB and are readily interpreted (Fig. 2, Supplementary Fig. 1)^22,31^. The slopes from these models are equivalent to AGB density (g m^-3^, calculated by dividing g m^-2^ by mean canopy height). Species-level densities ranged between 375 g m^-3^ to 13,801 g m^-3^ (Supplementary Fig. 1, Supplementary Table 2). Mean canopy height was an accurate predictor for individual species, especially when calibrated for specific ecophenotypic and phenological conditions^15,31,41^. Model goodness-of-fit was strong, with adjusted R^2^ values ranging from 0.46 to 0.99 with a mean of 0.83 (Supplementary Fig. 1, Supplementary Table 2). Leave-one-out cross-validation indicated a mean prediction error of 7.4% (Supplementary Table 2). The high goodness-of-fits indicated the photogrammetric approach performed as well as widely used *in situ* allometric approaches at the species-level (Fig. 1, Table 1, Supplementary Fig. 1, Supplementary Table 2)^15,22,41,42^. Importantly, however, intensive drone surveys are relatively easy to conduct over larger spatial extents of several hectares. Using a carefully designed, standardised protocol^33^ for acquiring and processing datasets yielded a good level of success in reconstructing 93% (688/741) of plots (Fig. 1C). The few instances where reconstructions were unsuccessful include mosses in rocky terrain, tall and dense grassland, and taller trees and shrubs (> 3 m) and are discussed in Supplementary Note 1. The similarities of the height-biomass relationships indicate this approach is generalisable across growth forms and environmental settings.

At the PFT-level, we found canopy height strongly predicted AGB across all six PFTs, with adjusted R^2^ ranging from 0.49 to 0.99 (Fig. 2, Table 1). For every 1-centimetre increase in mean canopy height, AGB increased by between 11 to 115 g m^-2^, depending on PFT (Fig. 2, Table 1). Ferns had the lowest density (1,096 g m^-3^), followed by forbs (1,191 g m^-3^), then graminoids (2,898 g m^-3^) and shrubs (3,214 g m^-3^) with notably similar densities, then small trees (5,572 g m^-3^) and lastly succulents had the largest density (11,532 g m^-3^). Species-level model slopes were generally similar *within*, but different *between*, PFTs. The similarity of densities within PFTs indicates these relationships are generally transferrable between species within PFTs, particularly for the better sampled types such as graminoids and shrubs, although phenotypic and phenological variation will limit accuracy^31,41^. Should destructive harvests for local calibrations not be possible due to resource limitations or taxon conservation status, the height-mass models described here could be used to non-destructively estimate AGB from similar drone-derived canopy height models (Table 1 and Supplementary Table 2). These allometric relationships were linear across the range of canopy height and biomass that we sampled, allowing their application from the whole plant-level to the ecosystem-level without necessarily requiring the discrete analysis of individual plants that can be challenging in ecosystems with coalesced canopies^14,27,31,43^.

Wind speed negatively affected canopy heights reconstructed from photogrammetry (Fig. 3A, Supplementary Table 3, Supplementary Fig. 2, Supplementary Fig. 3, Supplementary Note 2). We found the height-wind interaction parameter was strongly positive and highly significant (*p* < 0.0001) (Fig. 3A Supplementary, Table 3). This influence was seen at both the PFT-level (Supplementary Fig. 2A) and species-level (Supplementary Fig. 3). Biomass divided by height increased for surveys conducted in windier conditions, because the movement of foliage meant lower mean canopy heights were reconstructed from images that were acquired non-concurrently (see Supplementary Note 2 for extended discussion). However, wind effects had only limited influence in our study because most of our plots were surveyed in relatively light wind conditions (of < 4 m s^-1^) (Supplementary Fig. 2A). We expect sensitivity to wind speed differs between species because the effects of wind on foliage motion depend on canopy architecture and mechanical properties like limb stiffness^44^ (Supplementary Fig. 3, Supplementary Note 2). Previous studies in forest settings have reported contradictory effects of wind speed on canopy reconstructions^30,35^, but we think that these differences are linked with the spatial grain of analysis. Our study demonstrates the need to control for the influence of wind speed in future work particularly when surveying low-stature plant canopies.

Sun elevation had no effect on allometric density and by extension reconstructed plant height (Fig. 3B, Supplementary Fig. 2B, Supplementary Fig. 5, Supplementary Table 5). Cloudy conditions appeared to have a notable effect on allometric density; however, the imbalance in observations under cloudy and clear conditions (n=80 and n=620, respectively), meant this effect was not considered reliable (Supplementary Table 4, Supplementary Fig. 4). As with wind, previous studies in forest settings reported contradictory effects of elevation on canopy reconstructions^30,35^. However, illumination conditions affect photogrammetry in complex ways^37,45^, with the influence of sun elevation depending on the distribution and intensity of shadows as well as the properties of the camera sensor and user choices during processing (see Supplementary Note 3 for extended discussion). When comparing findings regarding illumination effects, it is therefore necessary to consider the capabilities of the sensors and workflows employed and the structural complexities of the observed ecosystems. Our findings suggest that surveying under low wind speeds may be a higher priority than optimal illumination conditions for obtaining structural models of vegetation in low stature ecosystems.

## Discussion

We established accurate height-biomass relationships for non-forest vegetation using standardised drone photogrammetry protocols. Our findings enable observations that will provide new insights into ecosystem dynamics at previously understudied scales across non-forested ecosystems. Linear models have strong correspondence with observations at the species and PFT-levels across a diverse range of low-stature ecosystems and perform as well as conventional *in situ* allometric approaches reported in the literature (Table 1, Fig. 2, Supplementary Table 2 and Supplementary Fig. 1). The similarity of graminoid and shrub PFT relationships indicate these could be applied together to estimate AGB in mixed ecosystems, without the need to individually classify these taxa, although there will be cases where allometric functions will need to be calibrated locally (Supplementary Note 4). As mean canopy height is readily compared between taxa, ecosystems and observation approaches^14,22^, these linear allometric relationships are straightforward to interpret^46^ and can be easily integrated with landscape modelling frameworks. Drone photogrammetry is a relatively ‘low-cost’ (although see Supplementary Note 5) tool well suited for local-scale observation in non-forest ecosystems. The ease of surveying landscape scales of 1 to 10 ha is critical to advancing beyond existing *in situ* approaches and overcoming the gap between on-the-ground monitoring and the coarser grain of global-scale products derived from satellite-based remote sensing^27,31^. Accurate information at these intermediary scales is invaluable for validating models and testing the scaling of ecological relationships and biomass carbon estimates from plots to biomes^6^.

Addressing critical knowledge gaps in plant science with drone photogrammetry requires standardised protocols, such as those used here, because photogrammetry-derived models are sensitive to the ways in which data are collected^27,30,35–38^, processed^27,36–38^, and analysed^20,27–29^. These sensitivities can be more pronounced for subjects with complex texture, such as vegetation, and hinder comparisons between products obtained from different workflows. To date, what has been missing are systematic and reproducible demonstrations of how drone data can be used in real-world plant ecology research. Using standardised protocols, we established comparable height-biomass relationships for a wide range of low-stature plant species for the first time and enable investigation of how factors such as wind speed (Fig. 3A), illumination (Fig. 3B)^35^, or antecedent conditions^41^ may influence allometric approaches. We show that it is important to account for the effects of wind speed during photogrammetric surveys beyond simply considering the effects of wind on drone platforms. The most reproducible reconstructions will be obtained under ‘zero’ wind speeds^30,35,37^, but this is often not possible under real world operational conditions^31,45,47^. Our findings demonstrate that data will be most comparable when wind speeds are similar, but also that, where differences are unavoidable, it will be possible to derive corrections for how wind influences canopy reconstructions from drone photogrammetry. We call for the continued development of harmonised and community-based protocols to maximise knowledge gains and support cross-biome syntheses^31,33,34,48^.

Our findings show drone photogrammetry can yield informative canopy height models capable of detecting ecologically significant differences in AGB across a diverse range of low-stature ecosystems globally. Drones have considerable advantages as data collection platforms for ecological applications, including their relatively low cost, versatility in deployment allowing high temporal resolution monitoring, and capacity to record fine-grained and spatially explicit data^34,45,49^. Systematic and comparable observations of plant canopy structure and biomass are vital for calibrating and evaluating vegetation models and biomass products retrieved from globally-available remote sensing systems^1,32,50,51^. Drone data collection can broaden the scope of research and monitoring programmes to obtain more representative observations in vulnerable and understudied low-stature ecosystems. Photogrammetric approaches for monitoring canopy height and biomass provide novel tools that should be used more widely by the ecological research community to improve assessments of ecosystem change and global carbon budgets.

## Data availability

Data collected for this publication, including aerial images, marker and plot coordinates, and dry sample weights, as well as site and survey metadata, are available from the NERC Environmental Information Data Centre (DOI: <DATA DEPOSIT IN PROGRESS> - AVAILABLE ON REQUEST IF REQUIRED FOR REVIEW). Code for photogrammetric processing and statistical analysis is available at <https://github.com/AndrewCunliffe/Global-Drone-Allometry>.

## Acknowledgements

We acknowledge funding support from NERC (NE/R00062X/1) awarded to R.E.B., A.M.C., K.A., S.S., NERC (NE/M016323/1) awarded to I.H.M.-S., NERC GEF (NERC/GEF:1063 and 1069) to I.H.M.-S. and A.M.C., U.S. Geological Survey Land Change Science Program awarded to M.L.V., NSF Graduate Research Internship Program (GRIP) awarded to C.A.H., USDA McIntire-Stennis project awarded to J.W.K., University of Cape Town award to S.C.P., J.S. and M.D.C., USDA Agricultural Research Service, GESFIRE (AGL2013-48189-C2-1-R), FIRESEVES (AGL2017-86075-C2-1-R), FIRECYL (LE033U14) and SEFIRECYL (LE001P17) awarded to J.M.F.-G., L.C. and S.S.-S. FPU16/03070 awarded to J.M.F.-G., NSF (#1836861), NASA ABoVE award number: NNX17AC58A, EU Horizon 2020 grant No. 776681 (PHUSICOS), Czech University of Life Sciences to J.P.; Jornada LTER (NSF 1832194), Australian Research Council (LP180100741) to M.L., National Council For Science And Technology to L.A.M.-B., Deutsche Forschungsgemeinschaft DFG (MA 2549/6-1 and RI 835/24-1) to I.M. and M.K. and the NSW Department of Industry (NCRIS co-funding of Terrestrial Ecosystem Research Network – Cumberland Plain Supersite) to M.M.B and A.G. We thank the following entities for permission to conduct sampling on their properties: Kelly Young, Deer Canyon Preserve, Drie Kuilen Nature Reserve, Département des Eaux et Forêts Maroc, Idaho Department of Fish and Game, Jornada LTER and ARS (Study number 545), Leroy Humphries, Nature Conservation Agency of the Czech Republic, Penrith Whitewater Stadium, SDSU Research Foundation, Sevilleta National Wildlife Refuge, The Idaho Chapter of The Nature Conservancy, USDI Bureau of Land Management, Utqiaġvik Inupiat Corporation, Worcester County Council, Natural England, Yukon Government and Parks (Permit: Inu-02-16) and The Inuvialuit People. We thank the following people for their assistance with data collection: Aaron Sobel, Alex Boehm, Alex Traynor, Andrew Corrales, Barbara Pachler, Ben Porter, Ben Spectre, Bobby Mullen, C. Wade Ross, Chad Radford, Craig McNamara, Danielle Lara, Dominic Fawcett, Eleanor Walker, Eric Spangler, Gerald Griesebner, Gerardo Armendariz, Haydn Thomas, James Atkins, Jason Wong, John Anderson, John Godlee, John Smith, Jordan Johnston, Julio Ceniceros, Julius Anchang, Justin Van Zee, Karen Anderson, Kathryn Fuller, Lara Prihodk, Mariana Orejel, Martin Barnett, Nick Smith, Qiuyan Yu, Ross Bryant, Roxanne Chepsongol, Sandra Angers-Blondin, Santeri Lehtonen, Seth Hall, Stephanie Baker, Travis Whitehead, Ulrich Zangerl, William Gentry and Zachary Winston. Any use of trade, firm, or product names is for descriptive purposes only and does not imply endorsement by the U.S. Government. The authors acknowledge the support of the University of Exeter’s High-Performance Computing (HPC) facility in conducting this study, and Tim Hill and Toby Pennington for useful discussions.

## Author contributions

A.M.C. conceived the research idea, administered the project, curated the data, did the data visualisation and led the writing of the manuscript. A.M.C. and K.A. developed the experimental design. A.M.C., K.A., R.E.B., I.H.M.-S., M.M.B., P.E.C., M.D.C., J.M.F.-G., A.G., N.P.H., C.A.H., P.H., J.W.K., I.M., L.A.M.-B., S.C.P., J.P., S.S., J.S., S.A.V. and M.L.V. acquired funding. A.M.C., K.A., F.B., R.E.B., I.H.M.-S., T.A., M.M.B., L.C., P.E.C., M.D.C., M.S.E.-L., S.M.E., J.M.F.-G., A.G.F., K.G., B.M.G., A.G., N.P.H., M.S.H., S.H., C.A.H., P.H., W.J., J.W.K., M.K., S.K., M.B.L., I.M., M.E.M, C.D.M., D.M., L.A.M.-B., S.C.P., J.P., E.S.-A., K.J.S., D.S.-B., P.S., S.S., J.S., C.S., S.S.-S., S.A.V., M.L.V., F.V., M.W., H.W. and R.W. undertook the investigation. A.M.C. and H.A.G. developed the processing program. A.M.C., I.H.M.-S. and H.A.G. performed the analysis. All authors contributed to the final version of the manuscript.

## References

1. Duncanson, L. et al. The importance of consistent global forest aboveground biomass product validation. Surv Geophys 40, 979–999 (2019).

2. Liu, Y. Y. et al. Recent reversal in loss of global terrestrial biomass. Nature Climate Change 5, 470–474 (2015).

3. Ahlström, A. et al. The dominant role of semi-arid ecosystems in the trend and variability of the land CO2 sink. Science 348, 895–899 (2015).

4. Poulter, B. et al. Contribution of semi-arid ecosystems to interannual variability of the global carbon cycle. Nature 509, 600–603 (2014).

5. Asner, G. P., Elmore, A. J., Olander, L. P., Martin, R. E. & Harris, A. T. Grazing Systems, Ecosystem Responses, and Global Change. Annual Review of Environment and Resources 29, 261–299 (2004).

6. Myers-Smith, I. H. et al. Complexity revealed in the greening of the Arctic. Nat. Clim. Chang. 10, 106–117 (2020).

7. McNicol, I. M., Ryan, C. M. & Mitchard, E. T. A. Carbon losses from deforestation and widespread degradation offset by extensive growth in African woodlands. Nature Communications 9, 3045 (2018).

8. Sleeter, B. M. et al. Effects of contemporary land-use and land-cover change on the carbon balance of terrestrial ecosystems in the United States. Environ. Res. Lett. 13, 045006 (2018).

9. Meilhac, J., Deschamps, L., Maire, V., Flajoulot, S. & Litrico, I. Both selection and plasticity drive niche differentiation in experimental grasslands. Nat. Plants 6, 28–33 (2020).

10. Meilhac, J., Durand, J.-L., Beguier, V. & Litrico, I. Increasing the benefits of species diversity in multispecies temporary grasslands by increasing within-species diversity. Ann Bot 123, 891–900 (2019).

11. Myers-Smith, I. H. et al. Eighteen years of ecological monitoring reveals multiple lines of evidence for tundra vegetation change. Ecol. Monogr. 89, (2019).

12. Harper, A. B. et al. Land-use emissions play a critical role in land-based mitigation for Paris climate targets. Nat Commun 9, 1–13 (2018).

13. Griscom, B. W. et al. Natural climate solutions. PNAS 114, 11645–11650 (2017).

14. Bartsch, A. et al. Feasibility of tundra vegetation height retrieval from Sentinel-1 and Sentinel-2 data. Remote Sensing of Environment 237, 111515 (2020).

15. Huenneke, L. F., Clason, D. & Muldavin, E. Spatial heterogeneity in Chihuahuan Desert vegetation: implications for sampling methods in semi-arid ecosystems. Journal of Arid Environments 47, 257–270 (2001).

16. Shriver, R. K. Quantifying how short-term environmental variation leads to long-term demographic responses to climate change. Journal of Ecology 104, 65–78 (2016).

17. Schimel, D. et al. Observing terrestrial ecosystems and the carbon cycle from space. Glob Change Biol 21, 1762–1776 (2015).

18. Bendig, J. et al. Estimating Biomass of Barley Using Crop Surface Models (CSMs) Derived from UAV-Based RGB Imaging. Remote Sensing 6, 10395–10412 (2014).

19. Kröhnert, M. et al. Watching grass grow -a pilot study on the suitability of photogrammetric techniques for quantifying change in aboveground biomass in grassland experiments. in ISPRS-International Archives of the Photogrammetry, Remote Sensing and Spatial Information Sciences vol. XLII–2 539–542 (Copernicus GmbH, 2018).

20. Grüner, E., Astor, T. & Wachendorf, M. Biomass prediction of heterogeneous temperate grasslands using an SfM approach based on UAV imaging. Agronomy 9, 54 (2019).

21. Wijesingha, J., Moeckel, T., Hensgen, F. & Wachendorf, M. Evaluation of 3D point cloud-based models for the prediction of grassland biomass. International Journal of Applied Earth Observation and Geoinformation 78, 352–359 (2019).

22. Cunliffe, A. M., Assmann, J., Daskalova, G., Kerby, J. T. & Myers-Smith, I. H. Aboveground biomass corresponds strongly with drone-derived canopy height but weakly with greenness (NDVI) in a shrub tundra landscape. Environmental Research Letters (2020) doi:10.1088/1748-9326/aba470.

23. Greaves, H. E. Applying Lidar and High-Resolution Multispectral Imagery for Improved Quantification and Mapping of Tundra Vegetation Structure and Distribution in the Alaskan Arctic. (University of Idaho, 2017).

24. Dubayah, R. et al. The Global Ecosystem Dynamics Investigation: high-resolution laser ranging of the Earth’s forests and topography. Science of Remote Sensing 100002 (2020) doi:10.1016/j.srs.2020.100002.

25. Westoby, M. J., Brasington, J., Glasser, N. F., Hambrey, M. J. & Reynolds, J. M. ‘structure-from-motion’ photogrammetry: A low-cost, effective tool for geoscience applications. Geomorphology 179, 300–314 (2012).

26. Dandois, J. P. & Ellis, E. C. High spatial resolution three-dimensional mapping of vegetation spectral dynamics using computer vision. Remote Sensing of Environment 136, 259–276 (2013).

27. Cunliffe, A. M., Brazier, R. E. & Anderson, K. Ultra-fine grain landscape-scale quantification of dryland vegetation structure with drone-acquired structure-from-motion photogrammetry. Remote Sens. Environ. 183, 129–143 (2016).

28. Wallace, L., Hillman, S., Reinke, K. & Hally, B. Non-destructive estimation of aboveground surface and near-surface biomass using 3D terrestrial remote sensing techniques. Methods in Ecology and Evolution 8, 1607–1616 (2017).

29. Lussem, U. et al. Estimating biomass in temperate grassland with high resolution canopy surface models from UAV-based RGB images and vegetation indices. JARS 13, 034525 (2019).

30. Frey, J. et al. UAV Photogrammetry of Forests as a Vulnerable Process. A Sensitivity Analysis for a Structure from Motion RGB-Image Pipeline. Remote Sensing 10, 912 (2018).

31. Poley, L. & McDermid, G. A systematic review of the factors influencing the estimation of vegetation aboveground biomass using unmanned aerial systems. Remote Sensing 12, 1052 (2020).

32. Bouvet, A. et al. An above-ground biomass map of African savannahs and woodlands at 25m resolution derived from ALOS PALSAR. Remote Sensing of Environment 206, 156–173 (2018).

33. Cunliffe, A. & Anderson, K. Measuring Above-ground Biomass with Drone Photogrammetry: Data Collection Protocol. Protocol Exchange (2019) doi:10.1038/protex.2018.134.

34. Tmušić, G. et al. Current Practices in UAS-based Environmental Monitoring. Remote Sensing 12, 1001 (2020).

35. Dandois, J. P., Olano, M. & Ellis, E. C. Optimal altitude, overlap, and weather conditions for computer vision UAV estimates of forest structure. Remote Sensing 7, 13895–13920 (2015).

36. James, M. R. & Robson, S. Mitigating systematic error in topographic models derived from UAV and ground-based image networks. Earth Surf. Process. Landforms 39, 1413–1420 (2014).

37. Mosbrucker, A. R., Major, J. J., Spicer, K. R. & Pitlick, J. Camera system considerations for geomorphic applications of SfM photogrammetry. Earth Surface Processes and Landforms 42, 969–986 (2017).

38. James, M. R., Antoniazza, G., Robson, S. & Lane, S. N. Mitigating systematic error in topographic models for geomorphic change detection: Accuracy, precision and considerations beyond off-nadir imagery. Earth Surface Processes and Landforms (2020) doi:10.1002/esp.4878.

39. Pätzig, M., Geiger, F., Rasche, D., Rauneker, P. & Eltner, A. Allometric relationships for selected macrophytes of kettle holes in northeast Germany as a basis for efficient biomass estimation using unmanned aerial systems (UAS). Aquatic Botany 162, 103202 (2020).

40. Whittaker, R. H. Communities and Ecosystems. (MacMillan Publishing Co, 1975).

41. Rudgers, J. A. et al. Sensitivity of dryland plant allometry to climate. Functional Ecology 32, 2290–2303 (2019).

42. Muldavin, E. H., Moore, D. I., Collins, S. L., Wetherill, K. R. & Lightfoot, D. C. Aboveground net primary production dynamics in a northern Chihuahuan Desert ecosystem. Oecologia 155, 123–132 (2008).

43. Krofcheck, D., Litvak, M., Lippitt, C. & Neuenschwander, A. Woody Biomass Estimation in a Southwestern U.S. Juniper Savanna Using LiDAR-Derived Clumped Tree Segmentation and Existing Allometries. Remote Sensing 8, 453 (2016).

44. Rowe, N. & Speck, T. Plant growth forms: an ecological and evolutionary perspective. New Phytologist 166, 61–72 (2005).

45. Aber, J. S., Marzolff, I, Ries, J. & Aber, S. W. Small Format Aerial Photography and UAS imagery: Principles, techniques and geoscience applications. (Elsevier, 2019).

46. Warton, D. I., Wright, I. J., Falster, D. S. & Westoby, M. Bivariate linefitting methods for allometry. Biological Reviews 259–291 (2006).

47. Duffy, J. P. et al. Location, location, location: considerations when using lightweight drones in challenging environments. Remote. Sens. Ecol. Conserv. 4, 7–19 (2017).

48. Pérez-Harguindeguy, N. et al. New handbook for standardised measurement of plant functional traits worldwide. Aust. J. Bot. 61, 167–234 (2013).

49. Anderson, K. & Gaston, K. J. Lightweight unmanned aerial vehicles (UAVs) will revolutionise spatial ecology. Frontiers in Ecology and Environment 11, 138–146 (2013).

50. Tian, F. et al. Remote sensing of vegetation dynamics in drylands: Evaluating vegetation optical depth (VOD) using AVHRR NDVI and in situ green biomass data over West African Sahel. Remote Sensing of Environment 177, 265–276 (2016).

51. Rodríguez-Fernández, N. J. et al. An evaluation of SMOS L-band vegetation optical depth (L-VOD) data sets: high sensitivity of L-VOD to above-ground biomass in Africa. Biogeosciences 15, 4627–4645 (2018).

52. Herold, M. et al. The role and need for space-based forest biomass-related measurements in environmental management and policy. Surv Geophys 40, 757–778 (2019).

53. Luhmann, T., Fraser, C. & Maas, H.-G. Sensor modelling and camera calibration for close-range photogrammetry. ISPRS Journal of Photogrammetry and Remote Sensing 115, 37–46 (2016).

54. James, M. R., Robson, S., d’Oleire-Oltmanns, S. & Niethammer, U. Optimising UAV topographic surveys processed with structure-from-motion: Ground control quality, quantity and bundle adjustment. Geomorphology 280, 51–66 (2017).

55. Hendrickx, H. et al. The reproducibility of SfM algorithms to produce detailed Digital Surface Models: the example of PhotoScan applied to a high-alpine rock glacier. Remote Sensing Letters 10, 11–20 (2019).

56. Nesbit, P. R. & Hugenholtz, C. H. Enhancing UAV-SfM 3D model accuracy in high-relief landscapes by incorporating oblique images. Remote Sensing 11, undefined-undefined (2019).

57. Chudley, T. R., Christoffersen, P., Doyle, S. H., Abellan, A. & Snooke, N. High-accuracy UAV photogrammetry of ice sheet dynamics with no ground control. The Cryosphere 13, 955–968 (2019).

58. Zhang, H. et al. Evaluating the potential of post-processing kinematic (PPK) georeferencing for UAV-based structure-from-motion (SfM) photogrammetry and surface change detection. Earth Surface Dynamics 7, 807–827 (2019).

59. Cunliffe, A. M. et al. Allometric relationships for predicting aboveground biomass and sapwood area of Oneseed Juniper (Juniperus monosperma) trees. Front. Plant Sci. 11, (2020).

60. Ribeiro-Gomes, K., Hernandez-Lopez, D., Ballesteros, R. & Moreno, M. A. Approximate georeferencing and automatic blurred image detection to reduce the costs of UAV use in environmental and agricultural applications. Biosystems Engineering 151, 308–327 (2016).

61. Wilke, N. et al. Quantifying lodging percentage and lodging severity using a UAV-based canopy height model combined with an objective threshold approach. Remote Sensing 11, 515 (2019).

62. PDAL Contributors. PDAL Point Data Abstraction Library. (2020).

63. R Core Team. R: A language and environment for statistical computing. (R Foundation for Statistical Computing, 2019).

64. Kennedy, S. Astral. (2020).

65. Stefan, V. plotbiomes: Plot Whittaker biomes with ggplot2. (2018).

66. Alfons, A. cvTools. (2015).

67. Bates, D., Mächler, M., Bolker, B. & Walker, S. Fitting Linear Mixed-Effects Models Using {lme4}. Journal of Statistical Software 67, 1–48 (2015).

68. Lüdecke, D. & Aust, F. ggeffects. (2020).

69. Krzywinski, M. & Altman, N. Error bars. Nature Methods 10, 921–922 (2013).

70. Kuznetsova, A., Brockhoff, P. B., Christensen, R. & Jensen, S. lmerTest. (2020).

71. Assmann, J. J., Kerby, J. T., Cunliffe, A. M. & Myers-Smith, I. H. Vegetation monitoring using multispectral sensors-best practices and lessons learned from high latitudes. Journal of Unmanned Vehicle Systems 334730 (2018) doi:10.1101/334730.

72. Lüdecke, D., Makowski, D., Waggoner, P. & Patil, I. performance. (2020).

