## Supplementary Materials for "Drone-derived canopy height predicts biomass across non-forest ecosystems globally"

**Cunliffe *et al*.**

This supplement contains:

1) Additional data visualisations of the species-level results (Supplementary Fig. 1),
2) Further information characterising the study sites and survey conditions (Supplementary Table 1),
3) Additional analysis and interpretation of how wind speed influences photogrammetric reconstructions of plants (Supplementary Figs. 2A and 3, Supplementary Table 3, Supplementary Note 2),
4) Additional analysis and interpretation of how cloud cover influences photogrammetric reconstructions of plants (Supplementary Fig. 4, Supplementary Tables 4 and 6),
5) Additional analysis and interpretation of how sun elevation influences photogrammetric reconstructions of plants (Supplementary Figs. 2B and 4, Supplementary Table 5, Supplementary Note 3),
6) Parameters for species-level height:biomass models (Supplementary Table 2),
7) Extended discussion of the limitations of photogrammetric reconstructions of plants (Supplementary Note 1, Supplementary Figs. 6 and 7),
8) Extended discussion of the limitations of universal allometric functions (Supplementary Note 4), and
9) Extended discussion of the true cost of photogrammetric surveys (Supplementary Note 5).

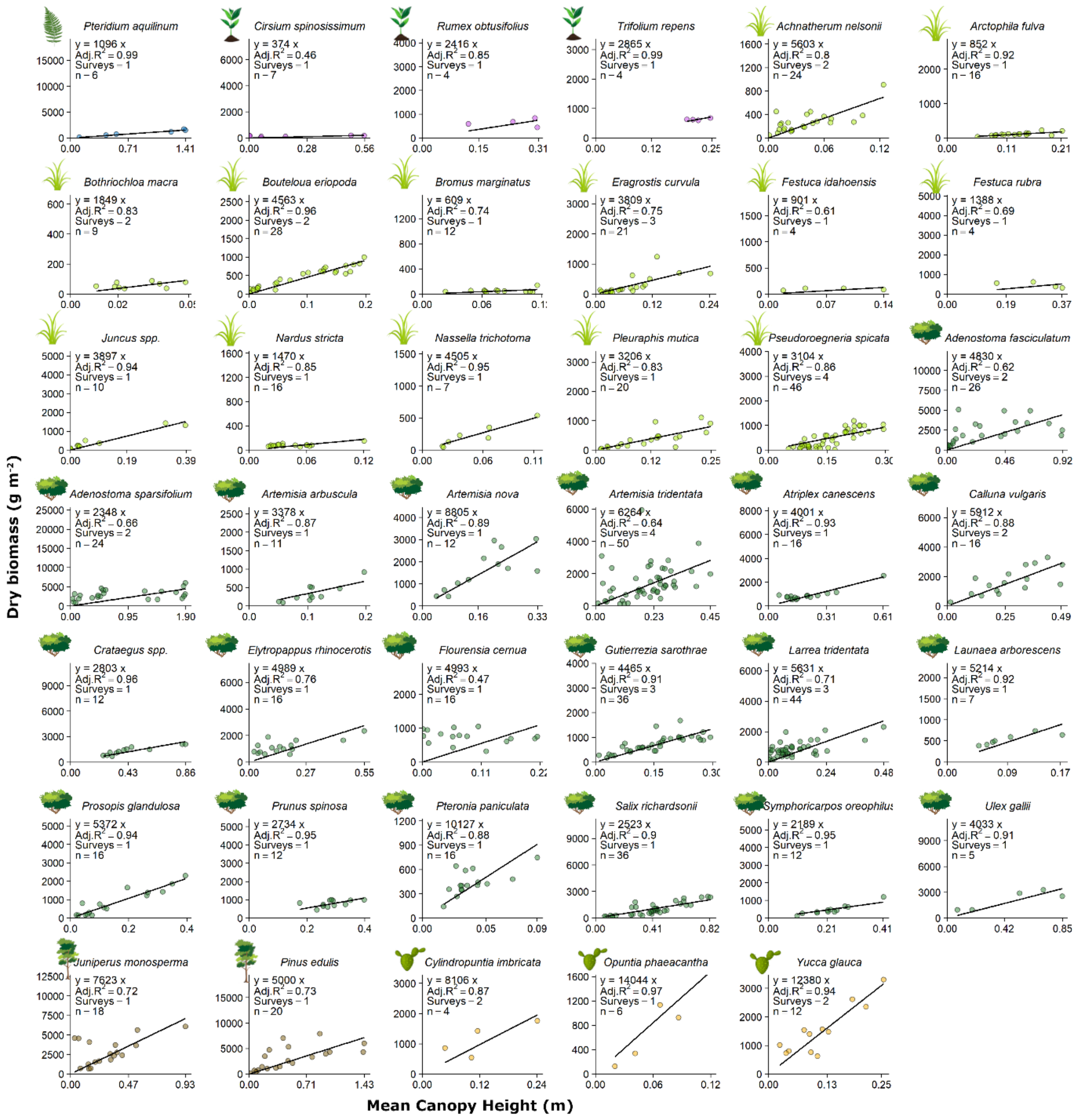


**Supplementary Fig. 1. Photogrammetrically-derived canopy height is a strong predictor of biomass across species.** We used ordinary least squares regression to fit linear models with an intercept constrained through the origin (solid black line) for all species with four or more observations. Species are grouped by plant functional type (indicated by icon and point colour). A constant X:Y ratio is used for all plots, so model slopes can be compared visually, even when axis ranges vary. Steeper slopes in these allometric models imply more biomass-per-unit-of-height, and model slopes are generally similar within plant functional groups. Full model results are included in Supp. Table 2.

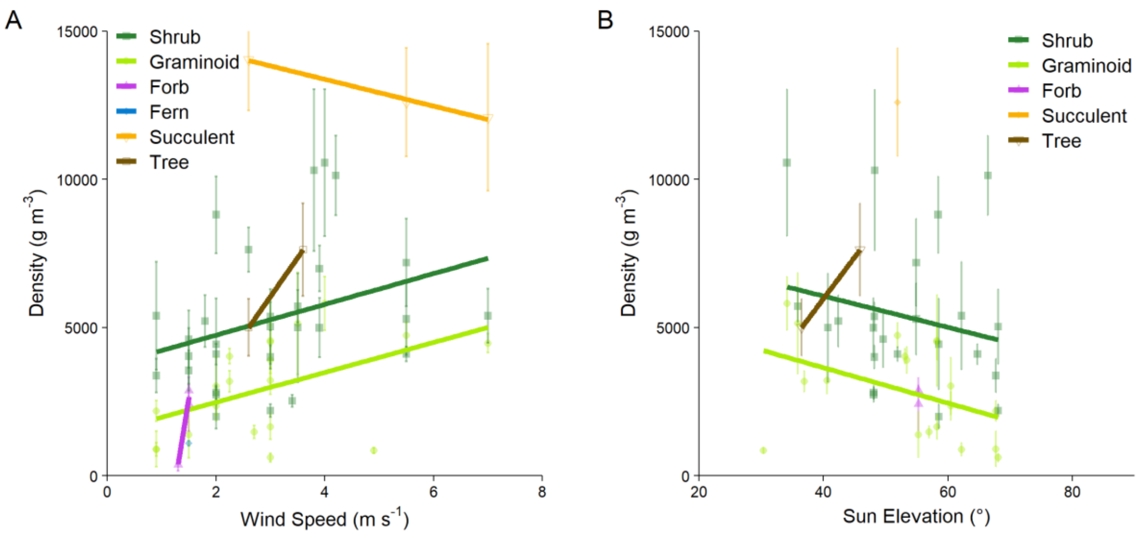


**Supplementary Fig. 2. Reconstructed plant height and thus height-biomass relationships were influenced by wind speed but were insensitive to sun elevation.** The allometric density (slope of height:biomass models ± 83% confidence intervals) calculated for each sample of each species plotted against wind speed (n=55) (A) and sun elevation for all surveys conducted under relatively clear sky conditions (n = 47, see methods for details) (B). Data are grouped by PFT, and linear models are fitted to illustrate the PFT-level trends tested with GLMMs (Supplementary Tables 3 and 4). We attribute the positive relationships between wind speed and density at the PFT-level (Fig. 3A) and species-level (Supplementary Fig. 3) to the influence of wind on reconstructed plant height (Supplementary Note 2). The negative relationships between sun elevation and density in the graminoid and shrub PFTs may be caused by lower sun angles causing shadowing that negatively bias reconstructed plant heights and thus increase density, but this effect was not statistically significant (Fig. 2B, Supplementary Table 5, Supplementary Note 3).

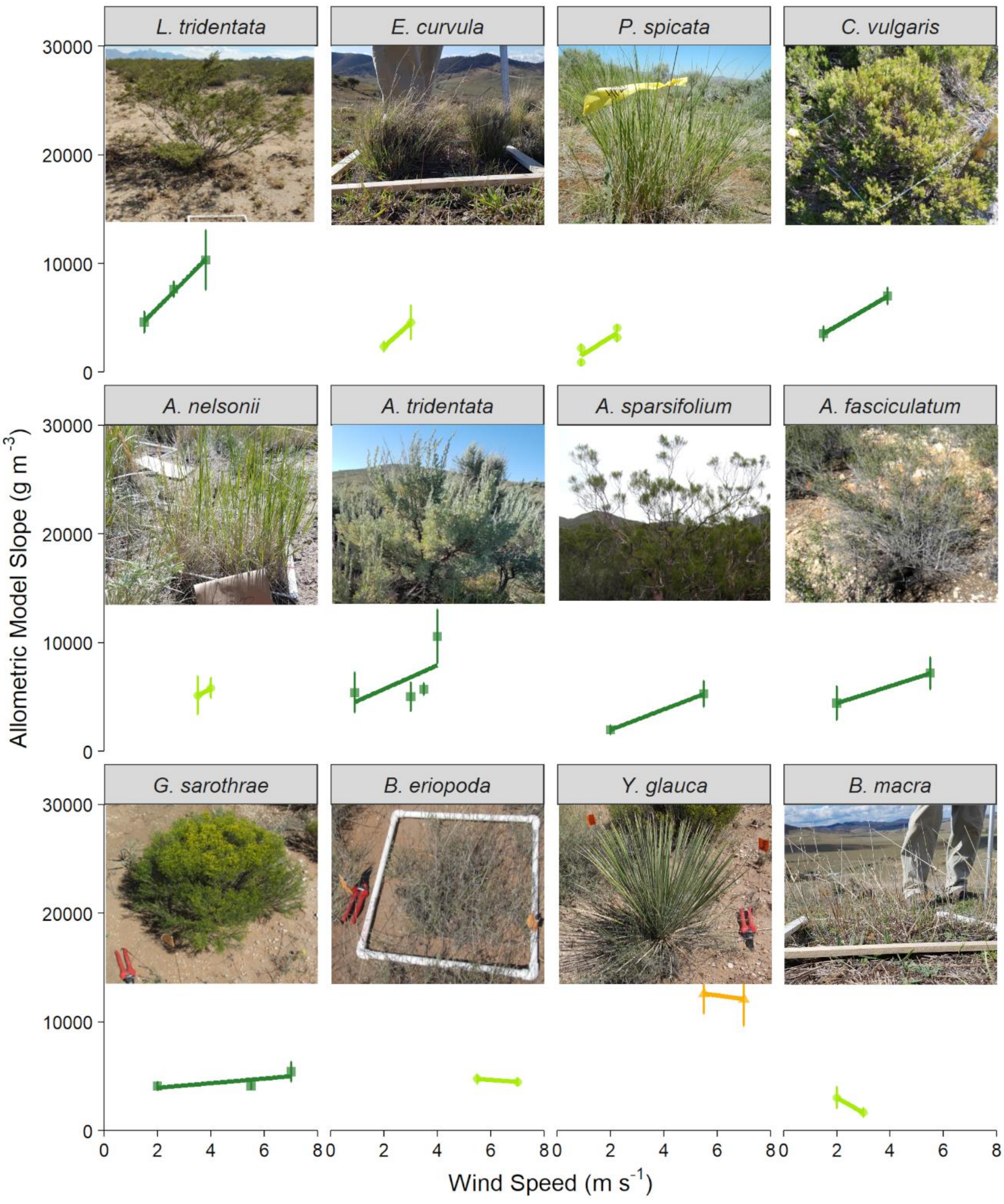


**Supplementary Fig. 3. Sensitivity of photogrammetrically-reconstructed height to wind speed differs between species based on growth form.** For the twelve species sampled more than once, the slope (± 83% confidence interval) of the linear model fitted to height and biomass for each sample was plotted against wind speed during the survey, and linear models were then fitted to these points to illustrate patterns at the species-level. Species are ordered by apparent sensitivity to wind speed, which broadly corresponded with canopy architecture. For further discussion see Supplementary Note 2.

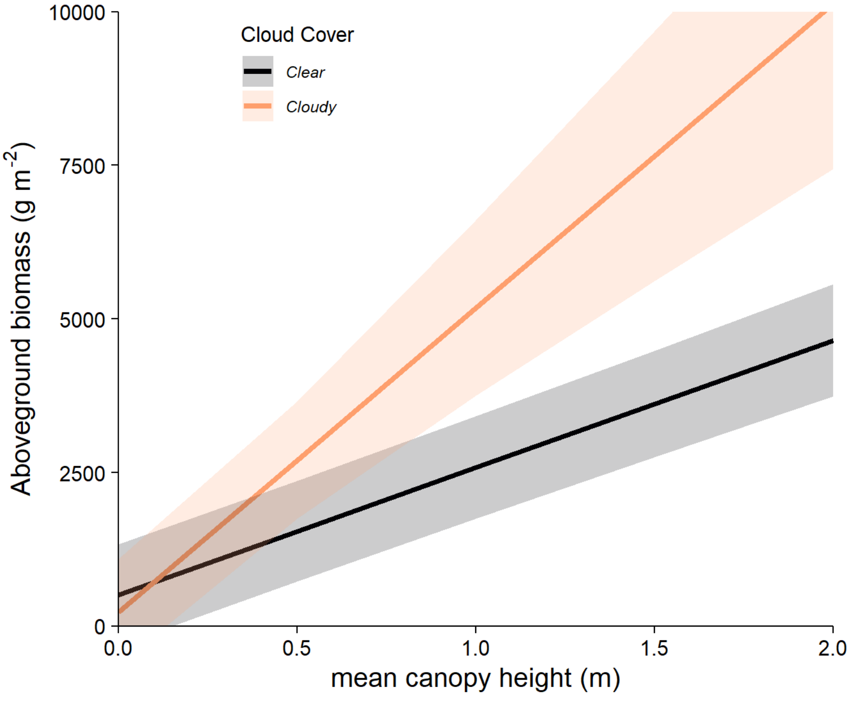


**Supplementary Fig. 4. The apparently strong effect of cloud cover on photogrammetrically-reconstructed height likely arises from imbalanced observations.** Mean predicted aboveground biomass variation over the range of observed mean canopy height. Shaded areas represent 95% confidence intervals on the model predictions. Cloud cover appears to strongly influence on the relationship between height and biomass; however, these results should be interpreted cautiously as these two factors are highly unbalanced in this analysis (‘Clear’ consisted of 620 observations from 33 surveys, whereas ‘Cloudy’ consisted of 80 observations from four surveys), and thus do not account for other possible covariates. Cloud cover had a statistically non-significant effect in the model, but there was a statistically significant interaction between cloud cover and height (Supplementary Table 4).

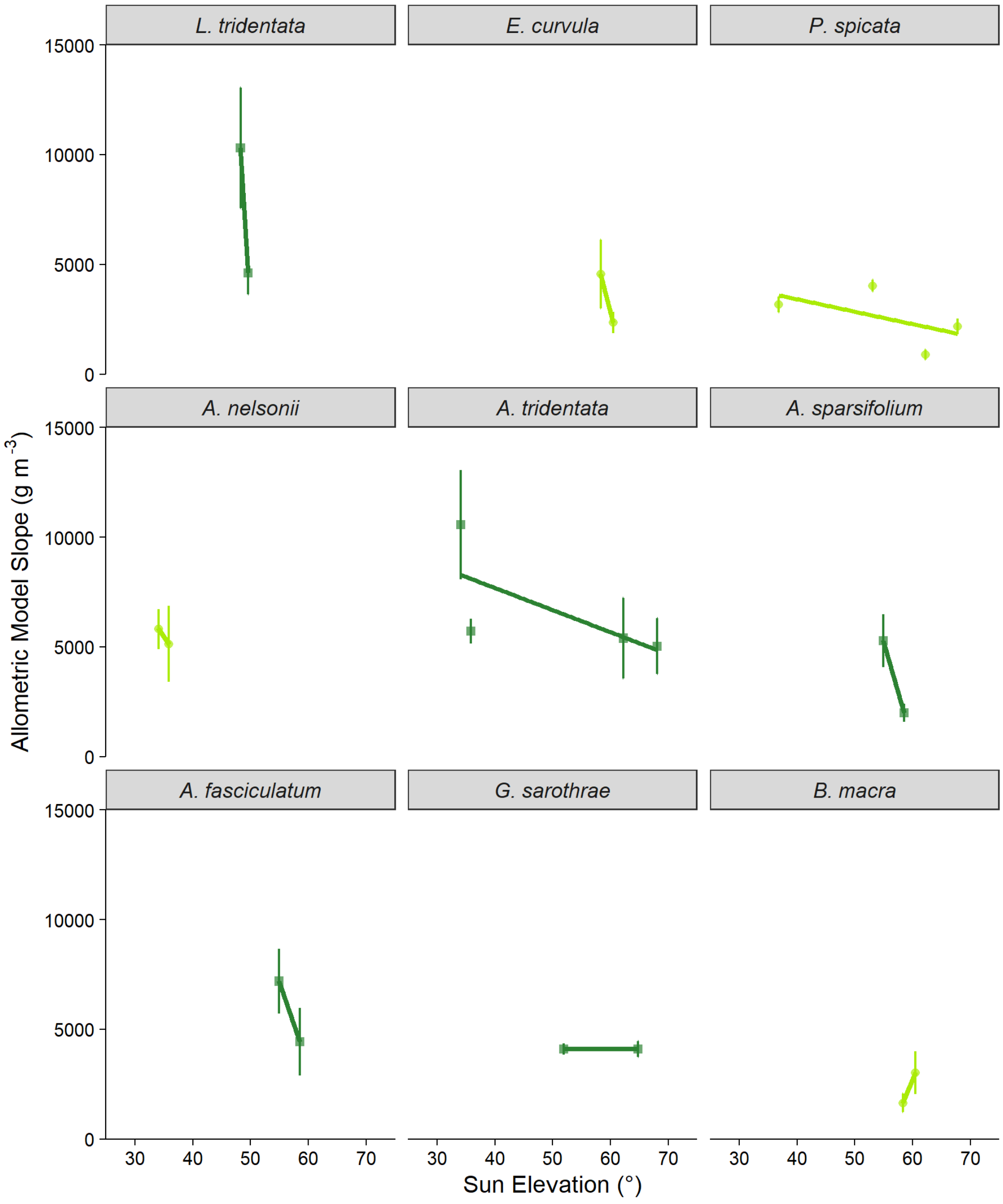


**Supplementary Fig. 5. Sun elevation has little systematic effect on photogrammetrically-reconstructed height at the species-level.** For the nine species sampled more than once under moderately clear skies (see methods for details), the slope (± 83% confidence interval) of the linear model fitted to height and biomass for each sample was plotted against sun elevation during the survey, and linear models were then fitted to these points to illustrate patterns at the species-level. For further discussion see Supplementary Note 3.

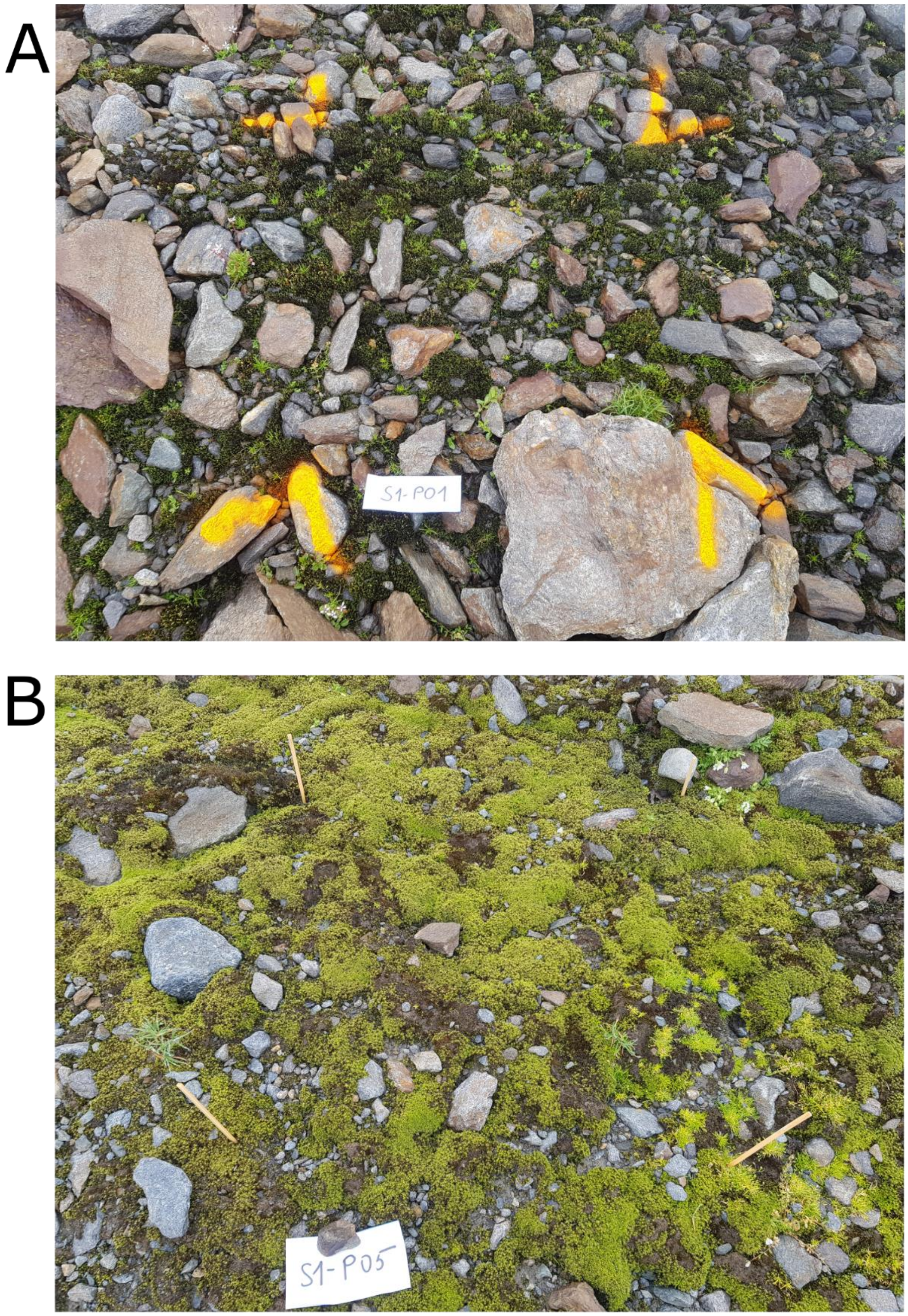


**Supplementary Fig. 6. This sampling approach was unable to usefully resolve the canopy height of mosses**. Photographs of two of the thirteen rocky bryophyte (moss) plots where we were unable to determine meaningful measurements of canopy heights due to the short height of the bryophytes (just a few centimetres) relative to the terrain roughness (**A** is of plot 20190810_HW_KS1_P01, and **B** is of plot 20190810_HW_KS1_P05). The 13 plots from these two sites were excluded from further analysis.

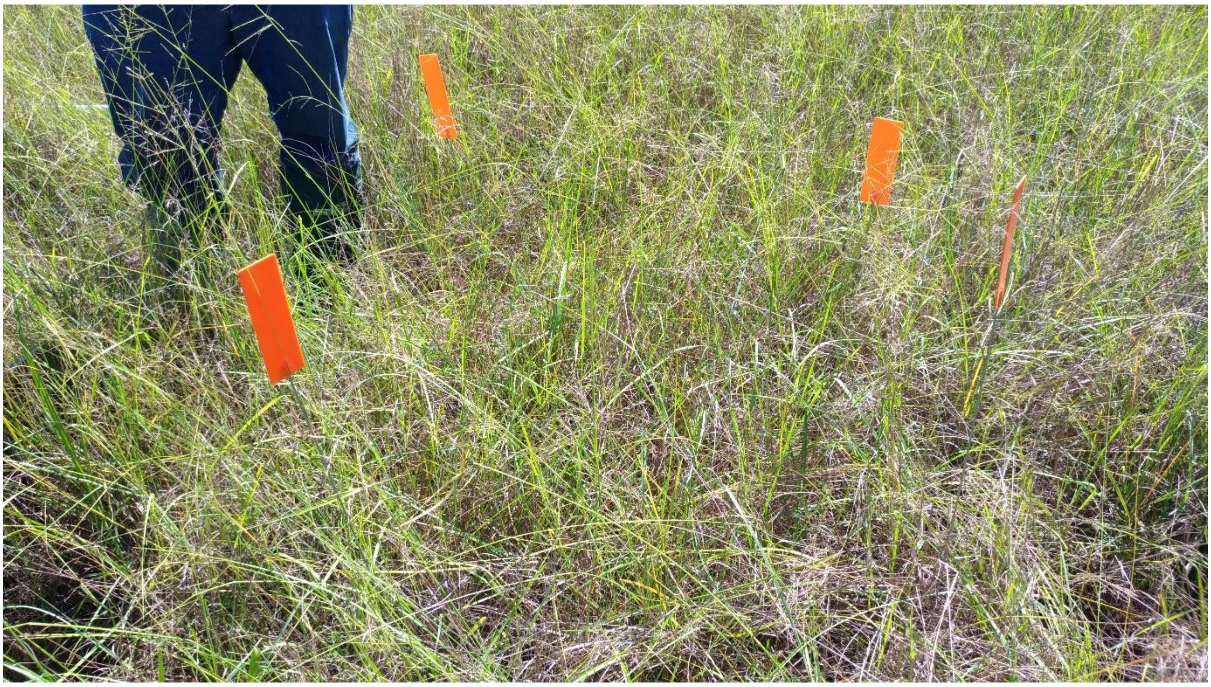

**Supplementary Fig. 7. Image alignment was not possible in this tall grassland, due to the complicated texture and structure of the subject preventing the accurate matching of tie points.** Photograph of a harvest plot (ID:20200313_AG_WSP_P01) in the one densely vegetated grassland that we were unable to reconstruct with the photogrammetry, co-dominated by *Eragrostis curvula, Chloris gayana* reaching up to 1.5 m in height. See Supplementary Note 1 for further details.

**Supplementary Table 1. Details of survey location, climate, ecosystem type and image sensor.** Where survey ID comprises the date of the survey (YYYYMMDD), followed by the initials of the lead contributor and three-character site code. MAT is mean annual temperature, MAP is mean annual precipitation, IGBP is the International Geosphere-Biosphere Programme code for the ecosystem type, and sky condition codes are after [1].

| Survey ID | Latitude | Longitude | Elevation (m) | MAT | MAP | Köppen class | IGBP | Wind speed (m s$^{-1}$) | Sky conditions | Drone & Sensor |
|---|---|---|---|---|---|---|---|---|---|---|
| 20160725_AC_ORC | 69.572 | -138.895 | 5 | -10.0 | 160 | ET | 6 | 3.4 | 5 | Tarot 680 Pro (Sony alpha6000 camera with 20 mm lens) |
| 20180602_MV_SGE | 37.979 | -109.936 | 1868 | 11.6 | 228 | BSk | 7 | 2 | 0 | 3DR Solo (Ricoh GR II camera) |
| 20180602_MV_SGB | 37.955 | -109.953 | 1855 | 11.6 | 228 | BSk | 7 | 2 | 0 | 3DR Solo (Ricoh GR II camera) |
| 20180623_JK_LCO | 43.357 | -114.393 | 1492 | 6.0 | 400 | BWh | 7 | 4 | 4 | DJI Phantom 4 Pro (FC6310 camera) |
| 20180624_JK_HAT | 43.420 | -114.411 | 1584 | 6.0 | 400 | BWh | 7 | 3.5 | 0 | DJI Phantom 4 Pro (FC6310 camera) |
| 20180626_FV_LAK | 52.251 | -2.254 | 35 | 10.5 | 606 | Cfb | 10 | 1.5 | 0 | DJI Phantom 4 Pro (FC6310 camera) |
| 20180727_JK_BC1 | 46.120 | -116.942 | 321 | 11.7 | 313 | Dfb | 10 | 2.25 | 0 | DJI Phantom 4 Pro (FC6310 camera) |
| 20180727_JK_BC2 | 46.127 | -116.941 | 279 | 11.7 | 313 | Dfb | 10 | 2.25 | 0 | DJI Phantom 4 Pro (FC6310 camera) |
| 20180812_FV_HBC | 52.333 | -2.259 | 53 | 10.5 | 606 | Cfb | 7 | 1.5 | 5 | DJI Phantom 4 Pro (FC6310 camera) |
| 20180915_AC_SEG | 34.362 | -106.702 | 1596 | 13.7 | 273 | BSk | 10 | 5.5 | 0 | DJI Phantom 4 Advanced (FC6310 camera) |
| 20180923_AC_SES | 34.335 | -106.745 | 1604 | 13.7 | 275 | BSk | 7 | 1.5 | 0 | DJI Phantom 4 Advanced (FC6310 camera) |
| 20181006_AC_WJS | 34.425 | -105.861 | 1931 | 11.5 | 364 | BSk | 9 | 3.6 | 4 | DJI Phantom 4 Advanced (FC6310 camera) |
| 20181022_SP_PTF | -33.695 | 19.895 | 875 | 14.8 | 372 | BSk | 7 | 3.9 | 2 | DJI Phantom 4 Advanced (FC6310 camera) |
| 20181023_SP_DKR | -33.571 | 20.029 | 1008 | 14.6 | 320 | BSk | 7 | 4.2 | 0 | DJI Phantom 4 Advanced (FC6310 camera) |
| 20190317_AC_SOC | 33.377 | -116.625 | 1415 | 13.6 | 533 | Csa | 6 | 5.5 | 2 | DJI Phantom 4 Advanced (FC6310 camera) |
| 20190326_AC_SO4 | 33.384 | -116.641 | 1447 | 13.5 | 554 | Csa | 6 | 2 | 3 | DJI Phantom 4 Advanced (FC6310 camera) |
| 20190605_PC_WBS | 43.167 | -116.715 | 1425 | 9.2 | 298 | Csa | 7 | 0.9 | 0 | DJI Phantom 4 (FC330 camera) |
| 20190617_JG_RLP | 43.044 | -5.413 | 1635 | 7.0 | 1600 | Cfb | 6 | 3.9 | 8 | DJI Phantom 4 Pro (FC6310 camera) |
| 20190625_PC_LOS | 43.143 | -116.738 | 1608 | 8.5 | 345 | Csa | 7 | 0.9 | 0 | DJI Phantom 4 (FC330 camera) |
| 20190626_TA_BRR | 50.478 | 9.971 | 870 | 4.8 | 1084 | Cfb | 10 | 2.7 | 0 | DJI Phantom 3 Advanced (FC300S camera) |
| 20190716_PC_MBS | 43.065 | -116.750 | 2100 | 5.4 | 803 | Csa | 7 | 3 | 0 | DJI Phantom 4 (FC330 camera) |

| | | | | | | | | | | |
|---|---|---|---|---|---|---|---|---|---|---|
| 20190807_SV_UTQ | 71.318 | -156.609 | 3 | -11 | 115 | ET | 11 | 4.9 | 4 | DJI Phantom 4 Pro (FC6310 camera) |
| 20190810_HW_KS1 | 46.861 | 10.731 | 2800 | 4.6 | 848 | ET | 16 | 1.3 | 8 | DJI Phantom 4 Pro (FC6310 camera) |
| 20190810_HW_KS2 | 46.864 | 10.733 | 2700 | 4.6 | 848 | ET | 16 | 1.3 | 8 | DJI Phantom 4 Pro (FC6310 camera) |
| 20190831_JP_DHR | 50.161 | 13.126 | 705 | 6.5 | 650 | Dfb | 7 | 2 | 4 | DJI Phantom 4 Pro (FC6310 camera) |
| 20191001_AC_SEG | 34.362 | -106.702 | 1596 | 13.7 | 273 | BSk | 10 | 7 | 7 | DJI Phantom 4 Advanced (FC6310 camera) |
| 20191003_AC_SES | 34.335 | -106.744 | 1604 | 13.72 | 275 | BSk | 7 | 2.6 | 7 | DJI Phantom 4 Advanced (FC6310 camera) |
| 20191011_AC_PIN | 34.439 | -106.234 | 2170 | 10.5 | 385 | BSk | 8 | 2.6 | 0 | DJI Phantom 4 Advanced (FC6310 camera) |
| 20191015_AC_JOC | 32.514 | -106.789 | 1300 | 14.6 | 240 | BSk | 7 | 3.8 | 4 | DJI Phantom 4 Advanced (FC6310 camera) |
| 20191015_AC_JOM | 32.610 | -106.796 | 1300 | 14.8 | 240 | BSk | 7 | 3 | 4 | DJI Phantom 4 Pro (FC6310 camera) |
| 20191015_AC_JOP | 32.667 | -106.770 | 1300 | 14.8 | 240 | BSk | 10 | 3 | 4 | DJI Phantom 4 Pro (FC6310 camera) |
| 20191015_AC_JOT | 32.514 | -106.740 | 1300 | 14.8 | 240 | BSk | 7 | 3.5 | 4 | DJI Phantom 4 Advanced (FC6310 camera) |
| 20191015_ML_CFT | -35.965 | 149.168 | 736 | 11.3 | 544 | Cfb | 10 | 2 | 4 | DJI Phantom 4 Pro (FC6310 camera) |
| 20191015_ML_OTH | -35.957 | 149.169 | 811 | 11.3 | 544 | Cfb | 10 | 3 | 4 | DJI Phantom 4 Pro (FC6310 camera) |
| 20191016_ML_RED | -35.968 | 149.164 | 731 | 11.3 | 544 | Cfb | 10 | 3 | 4 | DJI Phantom 4 Pro (FC6310 camera) |
| 20191024_SE_MEX | 27.251 | -110.192 | 4 | 23.7 | 317 | Bwh | 14 | 3 | 0 | 3DR Solo (MAPIR Survey 2 RGB camera) |
| 20200309_IM_IRG | 30.586 | -8.931 | 406 | 20.1 | 226 | BSh | 8 | 1.8 | 0 | DJI Phantom 4 Advanced (FC6310 camera) |
| 20200313_AG_WSP | -33.722 | 150.685 | 19 | 18.6 | 705 | Cfb | 10 | 1.6 | 0 | DJI Mavic Pro Platinum (FC220 camera) |

**Supplementary Table 2. Parameters for species-level linear models**, fitted for all species with four or more observations. Where PFT is plant functional type, n is number of observations, Adj. $R^2$ is the adjusted $R^2$, and LOOCV is the prediction error from Leave-One-Out Cross-Validation divided by the slope.

| PFT | Family | Species | n | Samples | Slope (g $m^{-2}$) | Residual standard error (g $m^{-2}$) | Adj. R2 | t-statistic | P value | LOOCV [%] |
|---|---|---|---|---|---|---|---|---|---|---|
| Fern | Dennstaedtiaceae | *Pteridium aquilinum* | 6 | 1 | 1096 | 53 | 0.99 | 20.558 | <0.0001 | 12.0 |
| Forb | Asteraceae | *Cirsium spinosissimum* | 7 | 1 | 374 | 140 | 0.46 | 2.660 | 0.0368 | 21.9 |
| Forb | Polygonaceae | *Rumex obtusifolius* | 4 | 1 | 2416 | 505 | 0.85 | 4.779 | 0.0174 | 8.9 |
| Forb | Fabaceae | *Trifolium repens* | 4 | 1 | 2865 | 112 | 0.99 | 25.435 | 0.0001 | 1.5 |
| Graminoid | Juncaceae | *Juncus spp.* | 10 | 1 | 3897 | 301 | 0.94 | 12.910 | <0.0001 | 3.3 |
| Graminoid | Poaceae | *Achnatherum nelsonii* | 24 | 2 | 5603 | 576 | 0.8 | 9.724 | <0.0001 | 1.4 |
| Graminoid | Poaceae | *Arctophila fulva* | 16 | 1 | 852 | 62 | 0.92 | 13.574 | <0.0001 | 1.3 |
| Graminoid | Poaceae | *Bothriochloa macra* | 9 | 2 | 1849 | 272 | 0.83 | 6.785 | 0.0001 | 1.0 |
| Graminoid | Poaceae | *Bouteloua eriopoda* | 28 | 2 | 4563 | 175 | 0.96 | 26.063 | <0.0001 | 1.6 |
| Graminoid | Poaceae | *Bromus marginatus* | 12 | 1 | 609 | 102 | 0.74 | 5.931 | 0.0001 | 2.8 |
| Graminoid | Poaceae | *Eragrostis curvula* | 21 | 3 | 3809 | 481 | 0.75 | 7.908 | <0.0001 | 1.8 |
| Graminoid | Poaceae | *Festuca idahoensis* | 4 | 1 | 901 | 335 | 0.61 | 2.684 | 0.0605 | 6.2 |
| Graminoid | Poaceae | *Festuca rubra* | 4 | 1 | 1388 | 437 | 0.69 | 3.175 | 0.0497 | 19.1 |
| Graminoid | Poaceae | *Nardus stricta* | 16 | 1 | 1470 | 153 | 0.85 | 9.554 | <0.0001 | 1.7 |
| Graminoid | Poaceae | *Nassella trichotoma* | 7 | 1 | 4505 | 393 | 0.95 | 11.458 | <0.0001 | 1.1 |
| Graminoid | Poaceae | *Pleuraphis mutica* | 20 | 1 | 3206 | 324 | 0.83 | 9.870 | <0.0001 | 2.2 |
| Graminoid | Poaceae | *Pseudoroegneria spicata* | 46 | 4 | 3104 | 187 | 0.86 | 16.569 | <0.0001 | 4.7 |
| Shrub | Asteraceae | *Artemisia arbuscula* | 11 | 1 | 3378 | 383 | 0.87 | 8.805 | <0.0001 | 3.7 |
| Shrub | Asteraceae | *Artemisia nova* | 12 | 1 | 8805 | 884 | 0.89 | 9.952 | <0.0001 | 3.5 |
| Shrub | Asteraceae | *Artemisia tridentata* | 50 | 4 | 6264 | 664 | 0.64 | 9.434 | <0.0001 | 7.4 |
| Shrub | Asteraceae | *Elytropappus rhinocerotis* | 16 | 1 | 4989 | 693 | 0.76 | 7.192 | <0.0001 | 8.1 |
| Shrub | Asteraceae | *Flourensia cernua* | 16 | 1 | 4993 | 1276 | 0.47 | 3.912 | 0.0014 | 8.4 |
| Shrub | Asteraceae | *Gutierrezia sarothrae* | 36 | 3 | 4465 | 230 | 0.91 | 19.344 | <0.0001 | 2.1 |

| | | | | | | | | | | |
|---|---|---|---|---|---|---|---|---|---|---|
| Shrub | Asteraceae | *Launaea arborescens* | 7 | 1 | 5214 | 559 | 0.92 | 9.320 | 0.0001 | 2.2 |
| Shrub | Asteraceae | *Pteronia paniculata* | 16 | 1 | 10127 | 932 | 0.88 | 10.864 | <0.0001 | 0.9 |
| Shrub | Chenopodiaceae | *Atriplex canescens* | 16 | 1 | 4001 | 276 | 0.93 | 14.465 | <0.0001 | 3.9 |
| Shrub | Caprifoliaceae | *Symphoricarpos oreophilus* | 12 | 1 | 2189 | 149 | 0.95 | 14.688 | <0.0001 | 3.1 |
| Shrub | Ericaceae | *Calluna vulgaris* | 16 | 2 | 5912 | 546 | 0.88 | 10.822 | <0.0001 | 7.3 |
| Shrub | Fabaceae | *Prosopis glandulosa* | 16 | 1 | 5372 | 330 | 0.94 | 16.254 | <0.0001 | 2.5 |
| Shrub | Fabaceae | *Ulex gallii* | 5 | 1 | 4033 | 561 | 0.91 | 7.180 | 0.0019 | 15.7 |
| Shrub | Salicaceae | *Salix richardsonii* | 36 | 1 | 2523 | 142 | 0.9 | 17.670 | <0.0001 | 11.1 |
| Shrub | Rosaceae | *Adenostoma fasciculatum* | 26 | 2 | 4830 | 728 | 0.62 | 6.626 | <0.0001 | 17.1 |
| Shrub | Rosaceae | *Adenostoma sparsifolium* | 24 | 2 | 2348 | 338 | 0.66 | 6.930 | <0.0001 | 60.1 |
| Shrub | Rosaceae | *Crataegus spp.* | 12 | 1 | 2803 | 158 | 0.96 | 17.716 | <0.0001 | 8.0 |
| Shrub | Rosaceae | *Prunus spinosa* | 12 | 1 | 2734 | 173 | 0.95 | 15.783 | <0.0001 | 4.9 |
| Shrub | Zygophyllaceae | *Larrea tridentata* | 44 | 3 | 5631 | 539 | 0.71 | 10.445 | <0.0001 | 5.7 |
| Succulent | Asparagaceae | *Yucca glauca* | 12 | 2 | 12380 | 899 | 0.94 | 13.763 | <0.0001 | 2.4 |
| Succulent | Cactaceae | *Cylindropuntia imbricata* | 4 | 2 | 8106 | 1527 | 0.87 | 5.308 | 0.013 | 5.9 |
| Succulent | Cactaceae | *Opuntia phaeacantha* | 6 | 1 | 14044 | 1073 | 0.97 | 13.083 | <0.0001 | 1.5 |
| Tree | Cupressaceae | *Juniperus monosperma* | 18 | 1 | 7623 | 1096 | 0.72 | 6.955 | <0.0001 | 8.4 |
| Tree | Pinaceae | *Pinus edulis* | 20 | 1 | 5000 | 671 | 0.73 | 7.446 | <0.0001 | 17.7 |

**Supplementary Table 3. Generalised linear mixed model parameters testing wind effects.** Model predicting aboveground biomass (g $m^{-2}$) as a function of mean canopy height (m), with wind speed (m $s^{-1}$) as an interaction effect and plant functional type as a random-effect. See Fig. 3A, Supplementary Fig. 2A, Supplementary Fig. 3 and Supplementary Note 2 for more information. Interactions are denoted by a colon (“:”).

*Model call: Biomass ~ Height × Wind + (1 | Plant Functional Type)*

| **Term** | **Estimate** | **Standard error** | **t-statistic** | **P value** |
|---|---|---|---|---|
| Intercept | 480.96 | 21.11 | 22.780 | <0.0001 |
| Height | 1327.65 | 22.89 | 58.013 | <0.0001 |
| Wind speed | 17.98 | 5.84 | 3.108 | 0.0019 |
| Height:Wind | 340.87 | 24.63 | 13.840 | <0.0001 |

**Supplementary Table 4. Linear mixed model parameters testing cloud cover effects.** Model predicting aboveground biomass (g $m^{-2}$) as a function of mean canopy height (m) and cloud cover (‘Clear’ = direct illumination, ‘Cloudy’ = sun obscured) as fixed effects and plant functional type as a random-effect. Imbalance in observations between the cloud factors means these results should be interpreted cautiously, see Supplementary Fig. 4 for more information. The PFT random effect explained 62% of variance in the model. Interactions are denoted by a colon (“:”).

*Model call: Biomass ~ Height × Cloud + (1 | Plant Functional Type)*

| **Term** | **Estimate** | **Standard error** | **t-statistic** | **P value** |
|---|---|---|---|---|
| Intercept | 511.81 | 419.72 | 1.219 | 0.277 |
| Height | 2069.33 | 123.02 | 16.821 | <0.001 |
| Cloudy | -284.88 | 158.86 | -1.793 | 0.073 |
| Height:Cloudy | 2880.92 | 715.84 | 4.025 | <0.001 |

**Supplementary Table 5. Linear mixed model parameters testing sun effects.** Model predicting aboveground biomass (g $m^{-2}$) as a function of mean canopy height (m) and sun elevation (degrees) as fixed effects and plant functional type as a random-effect. See Fig. 3B, Supplementary Fig. 2B, Supplementary Fig. 5 and Supplementary Note 4 for more information. The PFT random effect explained 59% of variance in the model. Interactions are denoted by a colon (":").

*Model call: Biomass ~ Height × Sun_elevation + (1 | Plant Functional Type)*

| **Term** | **Estimate** | **Standard error** | **t-statistic** | **P value** |
|---|---|---|---|---|
| Intercept | 583.96 | 468.18 | 1.247 | 0.248 |
| Height | 2939.14 | 770.76 | 3.813 | <0.001 |
| Sun_elevation | -0.21 | 4.06 | -0.053 | 0.958 |
| Height:Sun_elevation | -16.53 | 14.38 | -1.150 | 0.251 |

**Supplementary Table 6. Sky Codes for qualitative classification of cloud-related ambient light conditions.** From Assmann *et al.*[1] after NERC Field Spectroscopy Facility, Edinburgh, UK (2018) based on work by Milton *et al.*[2].

| **Sky code** | **Condition** |
|---|---|
| 0 | Clear sky |
| 1 | Haze |
| 2 | Thin cirrus, sun not obscured |
| 3 | Thin cirrus, sun obscured |
| 4 | Scattered cumulus, sun not obscured |
| 5 | Cumulus over most of sky, sun not obscured |
| 6 | Cumulus, sun obscured |
| 7 | Complete cumulus cover |
| 8 | Stratus, sun obscured |
| 9 | Drizzle |

**Supplementary Note 1. Notes on the limitations of photogrammetric reconstructions of plants.**

While our approach usually yielded highly plausible reconstructions of the plants within the harvest plots (Fig. 1C), in a few isolated instances the method did not work well. With a view to sharing our experience with the community (e.g. [3]), we describe these challenges here.

We observed that taller (>3 m maximum height) plants were more likely to be poorly reconstructed by the photogrammetry (e.g. *Juniperus monosperma* and *Pinus edulis*), which is illustrated by the negative bias in canopy height relative to the fitted model for a few of the plots with greater biomass in the Shrub and Tree panels of Fig 2. We attribute this poorer reconstruction primarily to excessive parallax in our low altitude flights that were optimised for shorter plants. Parallax is the effect whereby the position or direction of an object appears to differ when viewed from different positions[4]. To overcome this issue, we suggest using higher survey altitudes for taller plants, testing longer focal lengths to maintain ground sampling distance while reducing parallax.

We tested the approach on mosses (bryophytes in the genera *Racomitrium* and *Pohlia*); however, we were unable to resolve meaningful measurements of canopy height in a rocky pro-glacial montane setting because the mosses were too short (just a few centimetres in height) relative to the terrain roughness (Supplementary Fig. 6).

We were unable to reconstruct usable results from a survey of a tall (up to 1.5 m) and dense perennial grassland, co-dominated by *Eragrostis curvula* and *Chloris gayana* (Supplementary Fig. 7). This site had a thick standing layer of dry dead grass stalks below the green live biomass. The complicated texture across this site confounded tie point matching, hindering the accurate estimation of exterior (location and orientation) and interior (lens distortion) camera parameters during the bundle adjustment phase of the structure-from-motion processing. The resultant dense cloud was particularly noisy, with many clearly erroneous points distorting canopy height measurements. Consequently, the 16 mixed-grass plots from this site were excluded from the graminoid plant functional type analysis. The acquisition of more precise camera locations through real time kinematic (RTK) or post-processed kinematic (PPK) type systems on drone platforms would help by better constraining the estimation of exterior and interior camera parameters; however, such complex scenes are likely to remain challenging settings for photogrammetry approaches.

*Flourensia cernua* presented the main exception to the otherwise consistent pattern of mean canopy height being a good predictor of biomass (Supplementary Fig. 1). The shrubs were particularly poorly reconstructed in that survey, producing weak correspondence between mean height and biomass. An unknown factor appeared to have destabilised the bundle adjustment, but the cause of this remains unclear as the image data were high quality (well exposed, correctly focused, with high overlap and strong network geometry) and that shrubland was open with ca. 70% bare ground so tie points should have been largely stable. The wind speeds during that survey were moderate, at ca 3.5 m $s^{-1}$.

**Supplementary Note 2. Notes on how wind speed influences canopy heights**

Our analysis indicates that canopy height reconstructed from drone-acquired photographs is sensitive to wind speed (Fig. 3A, Supplementary Fig. 2, Supplementary Fig. 3, Supplementary Table 3). The estimate for the height-wind interaction parameter in the generalised linear mixed model was strongly positive and statistically significant ($p < 0.0001$), indicating that the relationship between aboveground biomass and canopy height gets stronger as wind speed increases (Supplementary Table 3; Fig 3; Supplementary Fig. 2).

Biomass divided by height increased for surveys conducted in windier conditions because the movement of foliage due to wind meant lower mean canopy heights were reconstructed from images that were acquired non-concurrently. These lower canopy heights then cause steeper slopes in the allometric relationship between mean canopy height and aboveground biomass. While our exploratory analysis cannot account for variation due to ecophenotype, phenology[5] or disturbance history, the consistency of responses across PFT-level (Fig. 3, Supplementary Fig. 2) and independently modelled species-levels (Supplementary Fig. 3) suggests that the overall results are robust. Dandois *et al.*[6] reported wind speed had no effect on reconstructed canopy height in a temperate deciduous forest but Frey *et al.*[7] reported wind speed does affect reconstructions of coniferous forest canopies. We think differences in sensitivity to wind are linked with the spatial grain of analysis (e.g. 1 $cm^{-2}$ versus 1 $m^{-2}$), in turn connected with differences in sensitivity to canopy structures.

We expect sensitivity to wind speed differs between species because the effects of wind on foliage (leaf and branch) motion depend on canopy architecture and mechanical properties like limb stiffness[8,9]. Tadrist *et al.*[9] found that foliage movement under wind was dominated at low velocity by high frequency, large amplitude, velocity independent individual leaf motions, and at high velocity by branch-induced, large scale, velocity-dependent motion. Taller plants are more prone to movement in the wind, and we observed greater wind influence on shrubs with more open branching structures (e.g. *A. sparsifolium, A. fasciculatum, L. tridentata* and *A. tridentata*), but less influence on shrubs with more compact growth forms (e.g. *G. sarothrae*) (Supplementary Fig. 3). In plants with a more compact growth form, wind-induced movement may simply substitute different foliage elements into the same space at different photographs. Of the three species exhibiting negative relationships between wind speed and slope of the allometric function, the allometric slopes for *B. eriopoda, Y. glauca* and *B. macra* all had overlapping 83% confidence intervals (Supplementary Fig. 3), so their differences are not statistically significant (at $p = 0.05$)[10]. Furthermore, one of the *B. macra* sites (Site: 'CTF'; Supplementary Table 1) experienced a fire disturbance 14-months prior to sampling which may contribute to the apparently anomalous relationship between height:biomass and wind (Supplementary Fig. 3).

Wind-induced movement of subjects between image acquisitions during drone surveys hinders their reconstruction from structure-from-motion multi-view stereopsis[11–13]. Non-stationary subjects will reduce the number of tie points that are matched correctly and often also increase the number of erroneously matched tie points, which together increase uncertainty in the estimation of external (location and orientation) and internal (lens distortion) camera parameters during the bundle adjustment. This degradation of parameter estimate quality will depend on the scene, as a greater proportion of tie points will remain stable in ecosystems with a large proportion of bare ground. The increased error in camera parameters combined with movement of the subject means that there is less coincidence between the depth maps calculated for each photograph, which results in fewer points being reconstructed by the multi-view stereopsis for moving vegetation[11,12], especially when depth filtering is applied as is normal practice in multi-view stereopsis[4,7,14–16]. The resulting dense point clouds contain fewer and more uncertain points, which are further processed to set any negative canopy heights to zero. Consequently, lower mean canopy heights are reconstructed when vegetation is moving due to wind (Fig. 3, Supplementary Fig. 2, Supplementary Fig. 3).

The force exerted by wind is non-linearly related to wind speed. When moving air is stopped by a surface, the dynamic energy in the moving air is transformed into pressure that acts on the surface as a force:

$$\mathbf{F = \tfrac{1}{2}\, \boldsymbol{p}\, v^2\, A}$$

Where F is in Newtons, p is the density of air in kg $m^{-3}$ (ca. 1.2 at sea level), v is velocity in m $s^{-1}$, and A is surface area in $m^2$. Future investigations into the influence of wind on vegetation reconstructions should test whether force, rather than wind speed, might be a better predictor of the influence on reconstructed canopy height. Frey *et al.*[7] suggested that the sensitivity of reconstructed forest canopies to wind speed depended on the ground sampling distance, with lower sensitivity at coarse spatial grains. Advancing understanding the interaction between the movement patterns of foliage and its reconstruction from non-concurrent photographs will need further empirical work.

**Supplementary Note 3: Notes on how sun elevation influences canopy heights**

Sun elevation had a very weakly negative, though statistically significant, effect on allometric density and by extension reconstructed plant height (estimate -7.67, std. error, 3.4, $p = 0.03$) (Supplementary Table 4, Fig. 3B). The weakly negative trend between sun elevation and density (Supplementary Fig. 2B) was seen in the relatively well-sampled graminoid and shrub PFTs, but there was little systematic pattern at the species level (Supplementary Fig. 4). At lower sun elevations, increased shadowing may slightly reduce reconstructed canopy heights[6] and thus increase biomass per unit of height. Reports on the effect of sun elevation on deciduous and coniferous forest canopy reconstructions have been contradictory[6,7]. However, forests are not always directly comparable to low-stature ecosystems because they differ in the distribution and intensity of shadows and illumination conditions that can have complex effects on photogrammetry[4,17]. Sensitivity to shadowing depends also on the camera’s dynamic range, i.e. its capacity to capture information in the brightest and darkest parts of the frame at the same time, as well as image formats (especially bit depth), camera settings and processing algorithms, which have all improved in recent years[4,7,17].

**Supplementary Note 4: Limitations on ‘universal’ allometries**

There may be limitations to transferability of the ‘universal’ allometric relationships described here. Previous studies have found strong power-law allometric relationships for diverse plant growth forms[18–20] and some consistency in allometric relationships over time[21]. While some studies have reported good allometric estimates of biomass at the plant functional group level[18,19,22], others found species-specific allometric models to perform best[20,23] or caution about overextending site-specific allometric functions[18]. In our study, all graminoids sampled have perennial life cycle strategies but there might be systematic differences between perennials and annual growth forms that hinder the transferability of a universal allometry. Future studies seeking to apply this approach to annuals should undertake sampling to calibrate their size – mass relationships as an expression of phenotypic plasticity. We also note that some species are known to adopt different growth forms in response to local environmental conditions and disturbance. For example, *B. gracilis* can shift from a bunch grass growth form to sod grass under elevated grazing pressure, as well as at higher elevations or farther north in its range[24,25]. Similarly, *Prosopis* species can vary from many-stemmed shrub forms to tree forms when growing on stream or river terraces with access to groundwater. The implications of such shifts, for species known to exhibit variable morphology, should be quantified before this photogrammetric approach should be used to test differences in canopy height or biomass between growth forms. Nonetheless, the overall approach of predicting biomass from mean canopy height has been shown to work, particularly when calibrated to phenotype, ecophenotype based on site conditions, phenophase based on antecedent condition and growth form based on disturbance or grazing pressure [18,19,21–23].

**Supplementary Note 5. Notes on costs**

The photogrammetric approaches tested in this study have often been described as ‘low cost’ since suitable image data can be collected with a drone and camera system costing ca. $1500 USD or less. However, such assertions are subjective depending on resource availability, as there can be additional costs for equipment to geolocate control points and for specialised hardware and software for data processing[3,26]. In some cases, it may be possible to mitigate these processing costs by using scalable web services and/or collaborations between data collectors and data processors (as in this project).

As photogrammetric reconstructions require spatial control that is accurate in relative (rather than absolute) terms, it would be possible to employ less expensive geolocation instruments (such as tacheometers costing ca. $300, theodolites or total stations). Furthermore, the cost of GNSS equipment (both on the ground and on the drone) is falling as technology progresses, lowering barriers to wider participation.

**References**


1. Assmann, J. J., Kerby, J. T., Cunliffe, A. M. & Myers-Smith, I. H. Vegetation monitoring using multispectral sensors - best practices and lessons learned from high latitudes. *Journal of Unmanned Vehicle Systems* 334730 (2018) doi:10.1101/334730.
2. Milton, E. J., Schaepman, M. E., Anderson, K., Kneubühler, M. & Fox, N. Progress in field spectroscopy. *Remote Sensing of Environment* **113**, S92–S109 (2009).
3. Tmušić, G. *et al.* Current Practices in UAS-based Environmental Monitoring. *Remote Sensing* **12**, 1001 (2020).
4. Aber, J. S., Marzolff, I, Ries, J. & Aber, S. W. *Small Format Aerial Photography and UAS imagery: Principles, techniques and geoscience applications*. (Elsevier, 2019).
5. Rudgers, J. A. *et al.* Sensitivity of dryland plant allometry to climate. *Functional Ecology* **32**, 2290–2303 (2019).
6. Dandois, J. P., Olano, M. & Ellis, E. C. Optimal altitude, overlap, and weather conditions for computer vision UAV estimates of forest structure. *Remote Sensing* **7**, 13895–13920 (2015).
7. Frey, J. *et al.* UAV Photogrammetry of Forests as a Vulnerable Process. A Sensitivity Analysis for a Structure from Motion RGB-Image Pipeline. *Remote Sensing* **10**, 912 (2018).
8. Rowe, N. & Speck, T. Plant growth forms: an ecological and evolutionary perspective. *New Phytologist* **166**, 61–72 (2005).
9. Tadrist, L. *et al.* Foliage motion under wind, from leaf flutter to branch buffeting. *Journal of The Royal Society Interface* **15**, 20180010 (2018).
10. Krzywinski, M. & Altman, N. Error bars. *Nature Methods* **10**, 921–922 (2013).
11. Walter, J., Edwards, J., McDonald, G. & Kuchel, H. Photogrammetry for the estimation of wheat biomass and harvest index. *Field Crops Research* **216**, 165–174 (2018).

12. Kattenborn, T., Sperlich, M., Bataua, K. & Koch, B. Automatic Single Tree Detection in Plantations using UAV-based Photogrammetric Point clouds. *Int. Arch. Photogramm. Remote Sens. Spatial Inf. Sci.* **XL–3**, 139–144 (2014).
13. Poley, L. & McDermid, G. A systematic review of the factors influencing the estimation of vegetation aboveground biomass using unmanned aerial systems. *Remote Sensing* **12**, 1052 (2020).
14. Agisoft. *Agisoft PhotoScan User Manual: Professional Edition, V 1.4*. (2018).
15. Cunliffe, A. M., Brazier, R. E. & Anderson, K. Ultra-fine grain landscape-scale quantification of dryland vegetation structure with drone-acquired structure-from-motion photogrammetry. *Remote Sens. Environ.* **183**, 129–143 (2016).
16. Lussem, U. *et al.* Estimating biomass in temperate grassland with high resolution canopy surface models from UAV-based RGB images and vegetation indices. *JARS* **13**, 034525 (2019).
17. Mosbrucker, A. R., Major, J. J., Spicer, K. R. & Pitlick, J. Camera system considerations for geomorphic applications of SfM photogrammetry. *Earth Surface Processes and Landforms* **42**, 969–986 (2017).
18. Nafus, A. M., McClaran, M. P., Archer, S. R. & Throop, H. L. Multispecies allometric models predict grass biomass in semidesert rangeland. *Rangeland Ecology & Management* **62**, 68–72 (2009).
19. Sanaei, A., Ali, A., Ahmadaali, K. & Jahantab, E. Generalized and species-specific prediction models for aboveground biomass in semi-steppe rangelands. *J Plant Ecol* **12**, 428–437 (2019).
20. Pottier, J. & Jabot, F. Non-destructive biomass estimation of herbaceous plant individuals: A transferable method between contrasted environments. *Ecological Indicators* **72**, 769–776 (2017).
21. Cleary, M. B., Pendall, E. & Ewers, B. E. Testing sagebrush allometric relationships across three fire chronosequences in Wyoming, USA. *Journal of Arid Environments* **72**, 285–301 (2008).

22. Oliveras, I. *et al.* Grass allometry and estimation of above-ground biomass in tropical alpine tussock grasslands. *Austral Ecology* **39**, 408–415 (2014).
23. Chieppa, J., Power, S. A., Tissue, D. T. & Nielsen, U. N. Allometric Estimates of Aboveground Biomass Using Cover and Height Are Improved by Increasing Specificity of Plant Functional Groups in Eastern Australian Rangelands. *Rangeland Ecology & Management* **73**, 375–383 (2020).
24. Wasser, C. H. *Ecology and Culture of Selected Species Useful in Revegetating Disturbed Lands in the West*. (Fish and Wildlife Service, U.S. Department of the Interior, 1982).
25. USDA. *Blue Grama Bouteloua gracilis (Willd. ex Kunth.) Lag. ex Griffiths*. 1–3 https://plants.usda.gov/plantguide/pdf/pg_bogr2.pdf (2007).
26. Glendell, M. *et al.* Testing the utility of structure from motion photogrammetry reconstructions using small unmanned aerial vehicles and ground photography to estimate the extent of upland soil erosion. *Earth Surf. Process. Landforms* **42**, 1860–1871 (2017).
